# Micro- and nanoplastics alter electrophysiological brain patterns and reshape human neurodevelopmental trajectories

**DOI:** 10.64898/2026.09.18.752710

**Authors:** Cassidy L. Poynter, Marta Sánchez-Carbonell, Rebekah L. Kendall, Marcus A. Garcia, Danilo Machado de Melo, Roxanne N. McPeck, Carol Mirita, Jared M. Brown, Matthew J. Campen, Andrij Holian, Alberto Cruz-Martín, Julio Aguado

## Abstract

Environmental microplastics and nanoplastics (MNPs) are emerging contaminants that accumulate in human tissues, including the brain, yet their mechanistic impact on the nervous system remains poorly understood. Converging postmortem evidence indicates increased MNP burden in brains from individuals with dementia, suggesting a potential link between plastic accumulation and disorders of the nervous system. However, whether MNPs directly contribute to brain aging and ensuing neurodegenerative processes, or disrupt neurodevelopment, is unknown. Here, we show that exposure to environmentally relevant MNPs induces cellular senescence and innate immune transcriptional programs in human brain organoids. Using region-specific cortical organoids, we demonstrate that synthetic polystyrene (PS), polyethylene terephthalate (PET), low-density polyethylene (LDPE), and high-density polyethylene (HDPE), as well as environmental ocean-derived MNPs (eMNPs) isolated from the coast of Hawaii, elicit a robust neuroinflammatory response. Transcriptomic analyses revealed activation of senescence-associated secretory phenotype (SASP) pathways. MNP exposure further disrupted neurodevelopmental trajectories, impairing neuroectodermal differentiation and redirecting lineage commitment toward mesoderm-like states, with concomitant enrichment of choroid plexus-like populations and a marked imbalance in neuronal and glial differentiation. These developmental alterations occurred in parallel with a pro-inflammatory and senescent microenvironment, suggesting coordinated disruption of developmental and aging programs. Importantly, high-density multielectrode array recordings revealed that MNP exposure altered electrophysiological activity and neuronal network dynamics, demonstrating that MNP-driven perturbations translate into dysfunctional neurological outcomes. Collectively, our findings identify MNPs as potent disruptors of human neurophysiology, linking plastic exposure to altered developmental trajectories, cellular senescence, neuroinflammation, and impaired neuronal network activity, providing mechanistic insights into how plastic accumulation may contribute to impaired brain dynamics.

## Introduction

Although plastic pollution has emerged as a global environmental concern^1^, the extent to which micro- and nanoplastics (MNPs) impact organ function remains poorly understood. Recent evidence shows the presence of MNPs across multiple human organs, including the liver, kidney, and notably the brain, where their accumulation appears especially pronounced^2,3^. Importantly, postmortem analyses of human brain tissues have revealed that individuals diagnosed with dementia harbor higher levels of MNPs compared to age-matched controls, suggesting a potential association between plastic burden and neurodegenerative pathology^2^. However, whether MNP accumulation directly contributes to cellular and brain dysfunction, or instead reflects a byproduct of disease progression, remains unknown.

A key biological process increasingly implicated in both brain aging and neurodegeneration is cellular senescence. Senescent cells accumulate in the central nervous system (CNS) during normal aging and are further enriched in pathological contexts, where they contribute to chronic inflammation through the senescence-associated secretory phenotype (SASP)^4^. In the brain, senescence has been linked to neuronal dysfunction, impaired regenerative capacity, and the progression of hallmark neurodegenerative features^5–7^, including amyloid-β deposition and tau pathology^8–10^. Despite these advances, whether environmental stressors such as MNPs can induce senescence in human neural tissues, and thereby promote aging-associated phenotypes, remains largely unexplored.

Modelling the effects of environmental toxicants in the human brain has historically been limited by the lack of physiologically relevant systems. Human induced pluripotent stem cell (iPSC)-derived brain organoids have emerged as a powerful platform to overcome these limitations, as they recapitulate key aspects of human brain development, cellular diversity, and regional specification in a three-dimensional context^11^. Region-specific organoids, including cortical and midbrain models, enable the interrogation of cell type- and circuit-specific responses to external perturbations^12^, providing a unique opportunity to study how environmental exposures impact human neurodevelopment and aging. Here, we leveraged human cortical and midbrain organoids to investigate the impact of MNP exposure on brain development and age-related trajectories. We exposed organoids to four ultraviolet (UV)-aged, commonly encountered plastic types - polystyrene (PS), polyethylene terephthalate (PET), low-density polyethylene (LDPE), and high-density polyethylene (HDPE) - as well as environmentally relevant, ocean-derived nanoplastics (eMNPs) isolated from the coast of Hawaii. Across all tested plastics, we found that MNP exposure induces a robust senescence response, accompanied by the activation of aging-associated transcriptional programs and a pro-inflammatory milieu. Functionally, electrophysiological profiling of mature midbrain organoids revealed that MNP exposure disrupts patterns of neuronal activity, demonstrating that these molecular and cellular alterations are accompanied by functional consequences in established human neural networks.

Importantly, beyond aging-related phenotypes, MNP exposure profoundly disrupts neurodevelopmental trajectories. We observed impaired ectodermal differentiation, coupled with a redirection of lineage commitment toward mesoderm-like states, and a marked alteration in neuro-specific differentiation. These changes occur in parallel with increased neuroinflammation and senescence, suggesting that MNPs simultaneously perturb developmental and aging pathways in human neural tissues.

Together, our findings identify MNPs as a previously underappreciated environmental driver of neurodevelopmental disruption, cellular senescence, and altered neural function in the human brain. By demonstrating that MNP exposure not only remodels developmental and aging-associated programs but also disrupts electrophysiological activity in mature neural tissue, these results provide mechanistic and functional insight into the potential link between plastic accumulation and neurodegenerative disease and highlight the need to further investigate the long-term neurological consequences of widespread environmental plastic exposure.

## Results

### sMNPs alter human neurodevelopmental programs

To investigate how acute and prolonged early life exposure to synthetic micro- and nanoplastics (sMNPs) influences human cortical brain development, we exposed human iPSC-derived cortical organoids (COs) to a combination of sMNPs previously found in human postmortem brain^2^ at physiologically relevant concentrations of 10 and 1,000 μg ml^-1^, beginning on day 6 of differentiation and maintained exposure throughout development (Supplementary Fig. 1d). We collected COs at 8 and 30 days post exposure (d.p.e.) for downstream analysis. The sMNP preparation consisted of polystyrene (PS), polyethylene terephthalate (PET), low-density polyethylene (LDPE) and high-density polyethylene (HDPE), whose identity and physicochemical properties were confirmed by bright-field microscopy, ATR-FTIR spectroscopy, and zeta potential measurements (Supplementary Fig. 1a-c).

COs showed no significantly altered growth throughout the 30-day exposure period (Supplementary Fig. 1e). Brightfield imaging revealed particle material accumulating within exposed organoids, with larger aggregates observed at the higher exposure concentration (Supplementary Fig. 1f). Consistently, polarized light microscopy (PLM) further confirmed birefringent sMNP particles within CO tissue following exposure (Fig. 1a).

**Figure 1:**
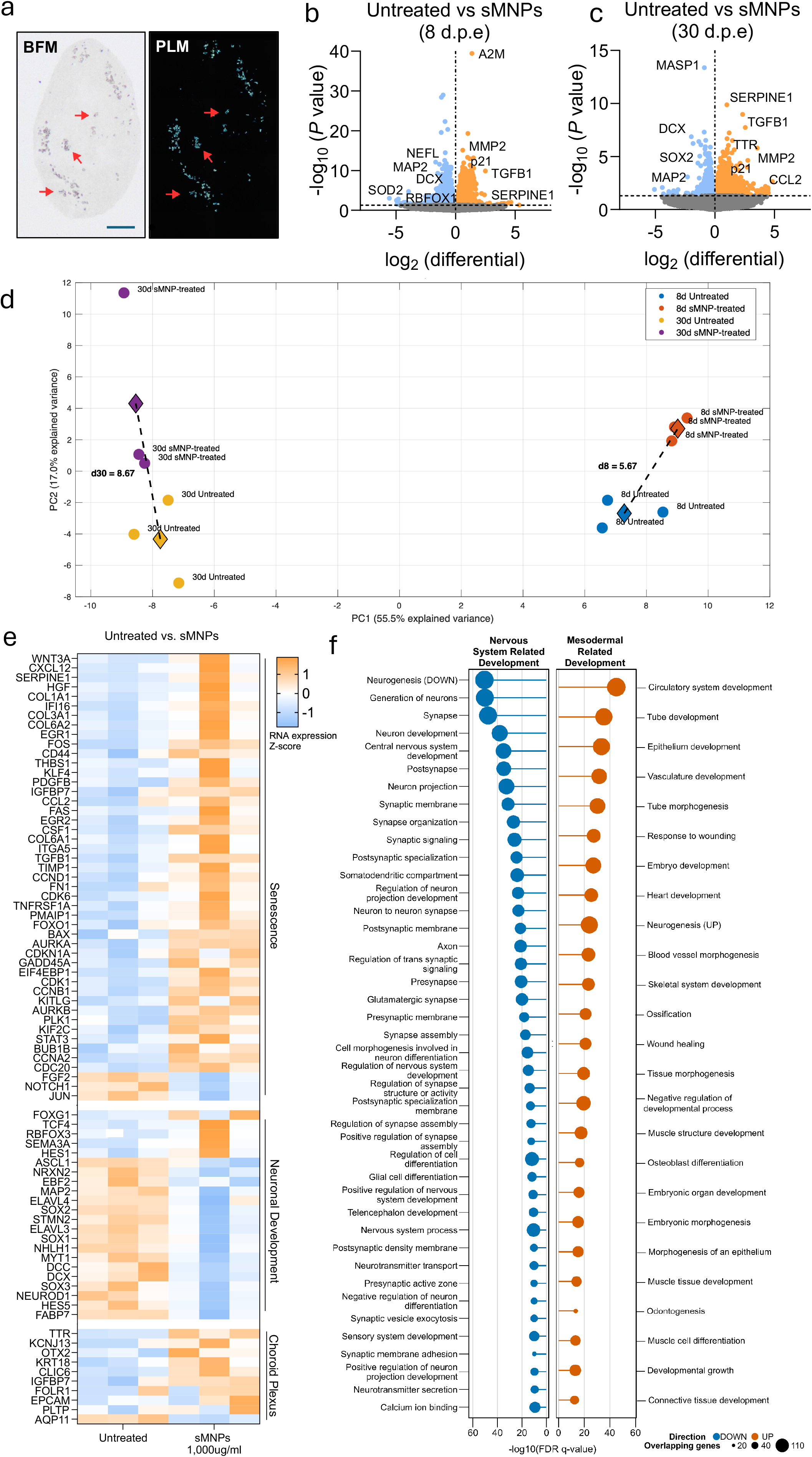
sMNP exposure suppresses neuronal developmental programs and promotes mesodermal and extracellular matrix remodeling in developing COs. (**a**), Representative bright-field microscopy (BFM; left) and polarized light microscopy (PLM; right) images of cortical organoids (COs) following 1,000 μg ml^-1^ synthetic micro- and nanoplastic (sMNP) treatment at day 6 of growth for 30 days. Red arrows indicate sMNP particles identified by their morphology in BFM and corresponding birefringence under PLM. Scale bar = 250 μm. (**b**) Volcano plots depicting differential gene expressions in untreated and 1,000 μg ml^-1^ sMNP-treated COs, exposed on day 6 of growth and collected 8 d.p.e. The x-axis shows log₂ fold change, and the y-axis shows −log_10_(*P* value). Significantly upregulated genes are shown in orange, and significantly downregulated genes are shown in blue; nonsignificant genes are shown in gray. Representative genes associated with neurodevelopment, senescence, and extracellular matrix remodeling are indicated. Significant thresholds were set at p<0.05. (**c**) Volcano plots depicting differential gene expressions in untreated and 1,000 μg ml^-1^ sMNP-treated COs, exposed on day 6 of growth and collected 30 d.p.e. The x-axis shows log_2_ fold change, and the y-axis shows −log10(*P* value). Significantly upregulated genes are shown in orange, and significantly downregulated genes are shown in blue; nonsignificant genes are shown in gray. Representative genes associated with neurodevelopment, senescence, and extracellular matrix remodeling are indicated. Significant thresholds were set at p<0.05. (**d**) Principal component analysis (PCA) of RNA-sequencing data in untreated and 1,000 μg ml^-1^ sMNP-treated COs, exposed on day 6 of growth and collected at 8 or 30 d.p.e. Each point represents an individual biological replicate, colored according to exposure and time point. PC1 (55.5% of explained variance) and PC2 (17.0% of explained variance) capture the major sources of transcriptomic variation. Diamonds represent group centroids, and dashed lines connect untreated and sMNP-treated centroids at each time point. Euclidean distances between centroids (D8 and D30) quantify the separation between treatment groups. (**e**) Heatmap showing the relative expression of selected genes in untreated and 1,000 μg ml^-1^ sMNP-treated COs, exposed on day 6 of growth and collected 30 d.p.e. Senescence-associated genes were identified from the SenSig^13^ and SenMayo^14^ gene sets. Gene expression values are displayed as row-wise Z-scores calculated from normalized RNA-sequencing expression data. (**f**) Gene Ontology (GO) enrichment analysis of significantly upregulated and downregulated differentially expressed genes (DEGs) in untreated and 1,000 μg ml^-1^ sMNP-treated COs exposed on day 6 of growth and collected 30 d.p.e. Enriched GO terms associated with downregulated DEGs are shown in blue and enriched GO terms associated with upregulated DEGs are shown in orange. Genes are ranked on p-value. Dot position represents enrichment significance (-log_10_(FDR q-value), and dot size corresponds to the number of genes associated with each GO term.

To determine whether sMNP exposure altered gene expression patterns, we performed bulk RNA sequencing following acute (8 d.p.e.) and prolonged (30 d.p.e.) sMNP exposure. We identified distinct transcriptional responses at both time points through differential expression analysis (Fig. 1b,c). Following acute exposure, we observed downregulated differentially expressed genes (DEGs) involved in neuronal development, including *NEFL*, *MAP2*, *RBFOX1* and *DCX* (Fig. 1b). Among upregulated DEGs, we found distinct regulators of senescence and cell cycle inhibitor genes such as *CDKN1A* (p21) and *SERPINE1* (Fig. 1b). By 30 d.p.e., we found this transcriptional profile sustained and further characterized by extracellular matrix remodeling and stress responses, including *TGFB1*, *MMP2*, and *CCL2* (Fig. 1c).

Using principal component analysis (PCA), we found component 1 as the primary source of transcriptomic variation related to age (PC1-55.5%), while component 2 explained the effects driven by sMNP exposure (PC2-17.0%) (Fig. 1d). Although untreated and sMNP-treated organoids clustered closely after 8 d.p.e., separation between treatment groups increased over prolonged exposure, indicating cumulative differences in developmental trajectories following early sMNP exposure.

To further characterize the transcriptomic changes induced by early life sMNP exposure, we examined curated gene sets associated with cellular senescence (SenSig^13^, SenMayo^14^), neurodevelopment, and choroid plexus identity. Transcriptomic analysis after acute exposure revealed altered expression levels of many genes associated with these signatures (Supplementary Fig. 1 g, h). Following prolonged exposure, we identified widespread upregulation of senescence-associated genes, whereas neurodevelopmental genes showed a downregulation in expression (Fig. 1e). Notably, we found enriched choroid plexus-associated genes (Fig. 1e), indicating significant developmental shifts induced by sMNP exposure in human cortical tissue, suggesting that both acute and prolonged sMNP exposure simultaneously activates stress-associated pathways while disrupting developmental programs required for normal forebrain maturation.

Next, we performed gene ontology (GO) enrichment analysis and demonstrated that downregulated DEGs from COs exposed to sMNP at 30 d.p.e. were enriched for processes involved in multiple neurological pathways, including nervous system development, neurogenesis, axonogenesis, synaptic organization, and neuronal differentiation (Fig. 1f). Furthermore, upregulated DEGs were enriched for mesoderm-related developmental processes, extracellular matrix organization, and tissue morphogenesis (Fig. 1f). These results collectively demonstrate that sMNP exposure in early development reprograms cortical transcriptomes by suppressing neurodevelopmental pathways and promoting senescence-associated and mesodermal transcriptional signatures.

To investigate the biological pathways altered following MNP exposure, we performed gene set enrichment analysis (GSEA) at 8 and 30 d.p.e. (Fig. 2a). Following acute exposure, we identified enrichment in pathways associated with cellular stress, inflammatory responses and aging; including mTORC1 signaling, TNFα signaling via NF-κB, interferon responses, p53 signaling, reactive oxygen species, and complement signaling. After prolonged exposure, we observed increased enrichment of pathways associated with cellular senescence and inflammation. We identified gene datasets such as cellular senescence, replicative senescence, and SenMayo (Fig. 2a), indicating a progressive shift from early cellular stress and inflammatory responses toward more pronounced senescence-associated signaling.

**Figure 2:**
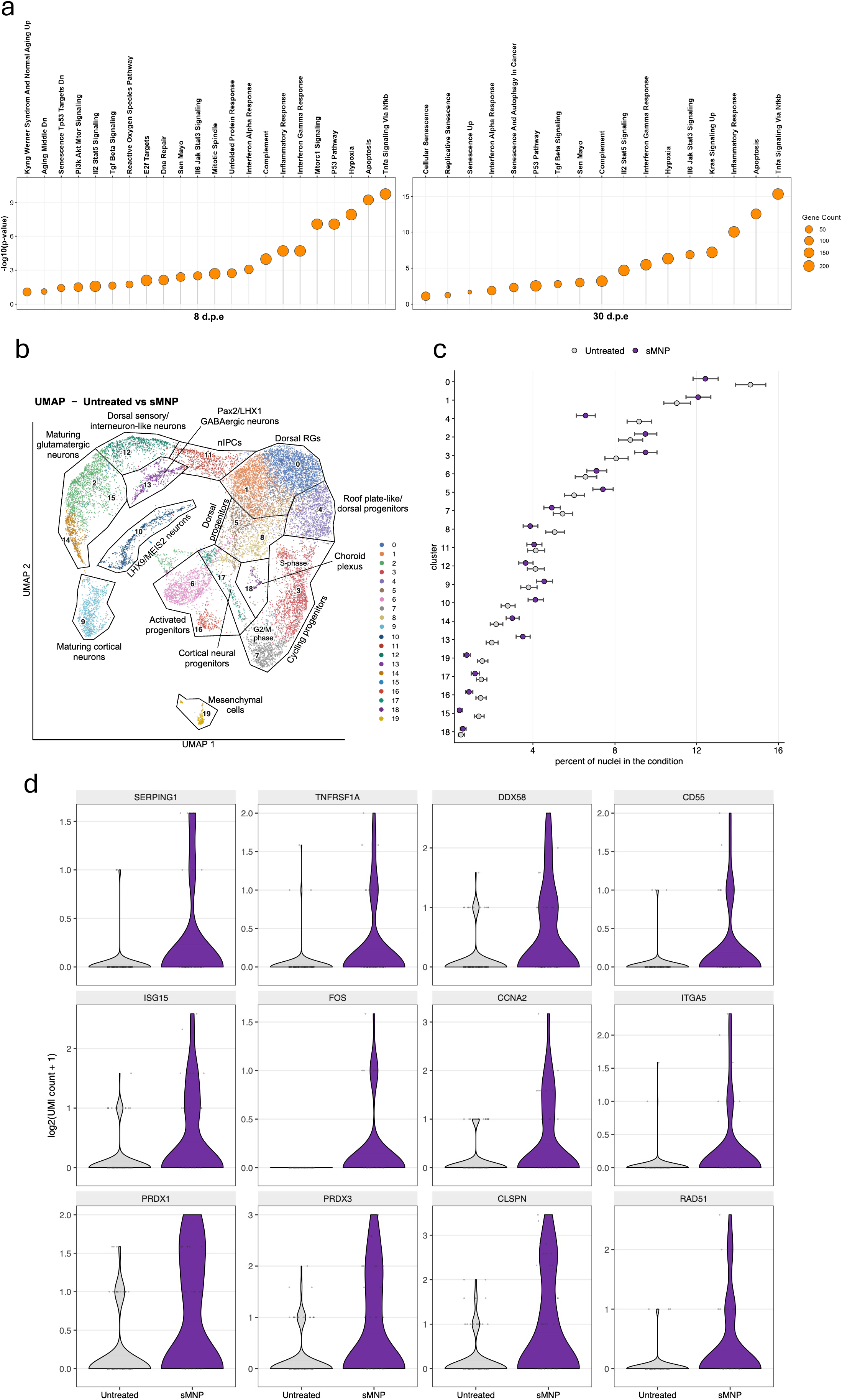
sMNP exposure remodels cortical development through cellular stress, inflammation, and altered neuronal states. (**a**) Gene set enrichment analysis (GSEA) of significantly upregulated pathways in untreated and 1,000 μg ml^-1^ sMNP-treated COs exposed on day 6 of growth and collected 8 d.p.e. and 30 d.p.e. Bubble plots display significantly enriched pathways ranked by statistical significance, represented as −log_10_(p-value), with bubble size corresponding to the number of genes contributing to each pathway. (**b**) UMAP visualization of single-nuclei RNA-sequencing data from untreated and 1,000 μg ml^-1^ sMNP-treated COs exposed on day 6 of growth and collected 30 d.p.e., showing the major transcriptionally defined cell populations identified across the dataset. Each point represents an individual nucleus, with colors indicating the corresponding cell cluster. Clusters were annotated based on established marker gene expression and include dorsal radial glia (RGs), neural intermediate progenitor cells (nIPCs), dorsal sensory/interneuron-like neurons, *Pax2/LHX1* GABAergic neurons, maturing glutamatergic neurons, *LHX6/MEIS2* neurons, cortical neural progenitors, activated progenitors, maturing cortical neurons, choroid plexus, roof plate-like/dorsal progenitors, cycling progenitors, and mesenchymal cells. (**c**) Relative composition of transcriptionally defined cell populations in untreated and 1,000 μg ml^-1^ sMNP-treated COs following 30 d.p.e. Data are presented as the proportion of nuclei assigned to each cell cluster within each treatment condition. Comparisons are based on relative proportions rather than raw cell counts because the total number of captured nuclei differed between conditions. Because each treatment condition was represented by a single 10x capture, differences in cell-type composition are presented as descriptive, hypothesis-generating observations and were not statistically tested for treatment effects. (**d**) Violin plots showing expression of representative differentially expressed genes in cluster 15, corresponding to maturing glutamatergic neurons, from untreated and 1000 μg ml^-1^ sMNP-treated COs exposed on day 6 of growth and collected 30 d.p.e. Genes represent functional programs associated with innate immune and inflammatory signaling (*SERPING1, TNFRSF1A, DDX58, CD55, ISG15*), oxidative stress (*PRDX1, PRDX3*), DNA damage and replication stress (*CLSPN, RAD51*), and stress-responsive, cell-cycle, and extracellular matrix signaling (*FOS, CCNA2, ITGA5*). Gene expression is shown as log_2_-transformed UMI counts + 1. Each violin represents the distribution of expression across cells within the indicated treatment group.

To infer the cellular identities associated with the transcriptional response to sMNP exposure, we deconvolved bulk RNA-seq signatures using a merged human single-cell reference derived from the Tabula Sapiens and Allen Brain Map datasets^15,16^. Upregulated and downregulated DEG signatures were analysed independently at both broad (Supplementary Fig. 2a) and high-resolution (Supplementary Fig. 3a) cell-type levels. Among genes upregulated following sMNP exposure, PCA revealed prominent vascular, mesenchymal and immune-associated transcriptional programs (Supplementary Figs. 2b–d and 3b–d). PC1 separated vascular/mesenchymal from immune-associated signatures, with endothelial and stromal populations enriched at one extreme and lymphoid and myeloid populations at the other (Supplementary Figs. 2c and 3c). PC2 further distinguished a mesenchymal/contractile program, predominantly mapping to contractile cells and fibroblasts, from a myeloid-enriched inflammatory program (Supplementary Figs. 2d and 3d). Thus, genes induced by sMNP exposure were preferentially associated with vascular, extracellular matrix and immune-related cellular programs.

**Figure 3:**
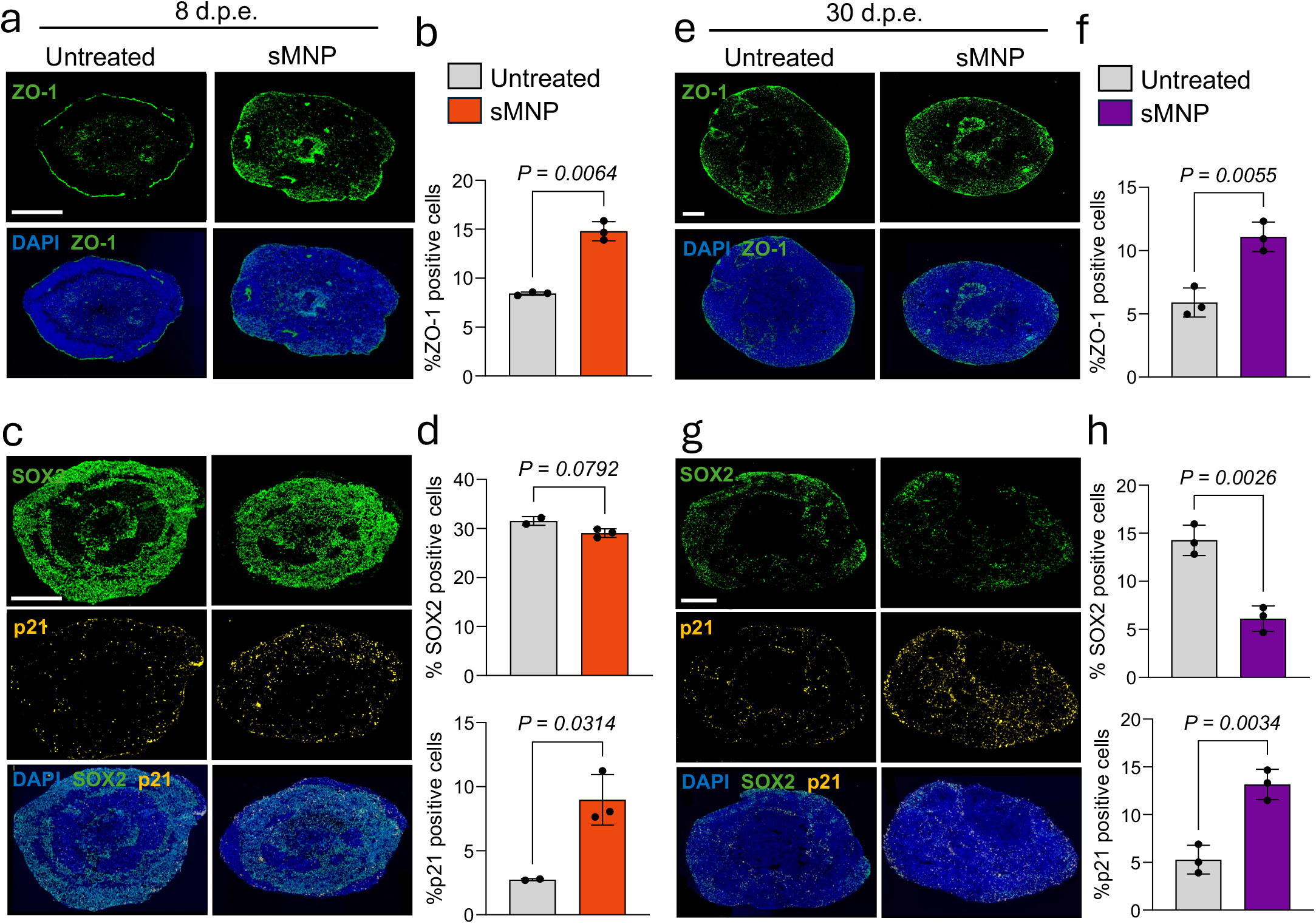
sMNP exposure drives time-dependent cellular stress, senescence, and neural progenitor dysfunction in developing COs. **a,c,e,g,** Representative immunofluorescence images of untreated and 1,000 μg ml^-1^ sMNP-treated COs exposed on day 6 of growth and collected at 8 d.p.e. (**a,c**) or 30 d.p.e. (**e,g**). Sections were stained for ZO-1 (green) (**a,e**); SOX2 (green) and p21 (yellow) (**c,g**). Nuclei were counterstained with DAPI (blue). Merged images are shown for each staining panel. Scale bars = 200 μm for ZO-1 and DAPI/ZO-1 images and 250 μm for SOX2, p21, and DAPI/SOX2/p21 images. **b,d,f,h,** Quantification of ZO-1-positive (**b,f**); SOX2 positive and p21-positive (**d,h**) cells in untreated and 1,000 μg ml^-1^ sMNP treated COs exposed on day 6 of growth and collected at 8 d.p.e. (**b,d**) or 30 d.p.e. (**f,h**). Data are presented as mean ± SD, with each dot representing an individual organoid. Statistical significance was determined using a two-tailed Welch’s *t*-test. Exact *P* values are shown.

Conversely, genes downregulated following sMNP exposure were strongly associated with neural populations (Supplementary Figs. 2e–g and 3e–g). PC1 was dominated by neuronal signatures, which at higher resolution mapped predominantly to excitatory intratelencephalic neurons, with additional contributions from interneuron and oligodendrocyte progenitor populations (Supplementary Figs. 2f and 3f). PC2 identified a distinct glial component dominated by oligodendrocyte-associated signatures (Supplementary Figs. 2g and 3g), consistent with a myelin-related co-expression module containing *PLP1, PMP2, ENPP2, MEGF10, SMOC1* and *PCDH15*. Collectively, these analyses associated sMNP-induced transcriptional changes with increased vascular, mesenchymal and immune-related programs and suppression of neuronal and glial programs.

To further resolve the cellular identities underlying the transcriptional changes identified by bulk RNA-seq, we performed single-nucleus RNA sequencing of untreated and sMNP-exposed cortical organoids following 30 days of exposure (Fig. 2b, c and Supplementary Fig. 7c). UMAP analysis resolved 20 transcriptionally distinct cell clusters spanning cortical radial glial and neural progenitor states, neurogenic intermediates, and immature-to-maturing neuronal populations, together with minor non-neural populations (Fig. 2b). sMNP exposure altered the relative representation of these developmental states (Fig. 2c). At the broader lineage level, progenitor and newborn-neuron populations significantly decreased from 45.9% to 42.4% (χ^2^(1) = 25.39, P < 0.0001), whereas immature neuronal populations significantly increased from 25.0% to 28.7% (χ^2^(1) = 35.04, P < 0.0001). These changes reflected coordinated shifts across multiple subpopulations, including reduced neurogenic radial glia and expansion of several immature neuronal states, along with more modest alterations in cycling progenitor populations (Fig. 2c). sMNP exposure was also accompanied by increased mesoderm- and extracellular matrix-associated transcriptional programs, consistent with the lineage-associated signatures identified by bulk RNA-seq.

We next examined transcriptional remodelling within specific neuronal populations and identified a pronounced response in maturing glutamatergic neurons (cluster 15) (Fig. 2d). sMNP exposure increased expression of genes associated with innate immune and inflammatory signalling (*SERPING1, TNFRSF1A, DDX58, CD55, ISG15*), oxidative stress (*PRDX1, PRDX3*), DNA damage/replication stress (*CLSPN, RAD51*), and stress-responsive, cell-cycle and extracellular matrix signalling (*FOS, CCNA2* and *ITGA5*, respectively) (Fig. 2d). These findings reveal a coordinated inflammatory and cellular stress response within maturing glutamatergic neurons, indicating that sMNP exposure alters not only the representation of cortical developmental states but also the molecular state of differentiating neurons.

Together with the bulk transcriptomic analyses, these findings indicate that prolonged sMNP exposure perturbs the developmental trajectory of human cortical organoids at both the transcriptional and cellular-state levels. Rather than causing a generalized loss of neuronal populations, sMNP exposure altered the balance between progenitor and neuronal states, suppressed normal neurodevelopmental programs, and promoted senescence-, inflammatory-, extracellular matrix-, and mesoderm-associated transcriptional signatures. These findings suggest that MNP exposure remodels cortical development through coordinated changes in cell-state composition and molecular identity.

### sMNP exposure alters cortical progenitor maintenance and senescence signaling

To determine whether these transcriptional changes were accompanied by alterations in cortical tissue organization, we assessed immunoreactivity of the tight junction marker, ZO-1 found in choroid plexus tissues; the neural progenitor marker, SOX2; and the cell cycle arrest and senescence marker, p21. At 8 d.p.e., sMNP-treated COs showed a significant increase in the percentage of ZO-1-positive regions compared with untreated organoids, indicating an early change of neuroepithelial tight junctions following exposure (Fig 3a, b). We observed a trend toward reduced SOX2-positive progenitor cells and a significant increase in p21 expression due to sMNP exposure, consistent with activation of a senescence-associated stress response (Fig 3c, d).

At 30 d.p.e., we observed persistent upregulation of ZO-1 in sMNP-treated organoids (Fig. 3 e, f). In addition, sMNP exposure significantly reduced SOX2-positive neural progenitors and increased p21-positive cells (Fig. 3 g, h). Together with the changes in inflammatory, cell cycle-related, developmental, and stress-response pathways identified by transcriptomic analysis (Fig. 1f and Fig. 2a), these data suggest that prolonged sMNP exposure alters neuroepithelial organization, reduces neural progenitor identity and activates a senescence-associated response.

To extend these observations, we assessed the cortical progenitor marker PAX6 and the senescence-associated marker p16. Consistent with the SOX2 analysis, immunofluorescence analysis revealed a significant reduction in PAX6-positive progenitor after acute exposure, whereas p16 expression remained unchanged (Supplementary Fig. 4a, c). By 30 d.p.e., PAX6 expression remained comparable between treatment groups, indicating distinct responses among progenitor markers (Supplementary Fig. 4b, d). SOX2 is broadly expressed in neuroepithelial stem cells and radial glia that maintain neural stem cell identity, whereas PAX6 is a transcription factor that specifies and maintains dorsal cortical progenitor fate^17,18^. Thus, the selective reduction in SOX2 may reflect impaired maintenance of early neural stem cell identity rather than a depletion of all cortical progenitor populations. The lack of a corresponding increase in p16 despite p21 induction further suggests that the senescence phenotype observed may be dispensable of the p16 senescence axis in developing COs.

### Individual sMNP polymers differentially alter cortical development

To determine whether individual sMNP polymers elicit early molecular changes during CO development, we exposed COs to PS, PET, LDPE, or HDPE, each at a concentration of 1,000 μg ml^-1^, beginning on day 6 of differentiation and collected organoids at 8 or 30 d.p.e. (Supplementary Fig. 5a). Although PS exposure modestly increased organoid size and HDPE exposure reduced size during the early exposure period, all organoids retained gross cortical morphology without structural abnormalities (Supplementary Fig. 5b). Using brightfield imaging, we further confirmed the persistent presence of sMNP particles within COs after both acute and prolonged exposure (Supplementary Fig. 5c, d).

DEG analysis revealed distinct transcriptional responses for each polymer compared with untreated controls (Fig. 4a). While the magnitude and composition of these responses varied among polymers, we identified several genes associated with extracellular matrix remodeling, neurodevelopment, and cellular stress that were commonly dysregulated across treatment groups. PS exposure produced greater alterations in extracellular matrix-associated genes, including *COL1A1* and *THBS2*, whereas PET exposure increased expression of genes involved in cellular stress and extracellular matrix signaling, such as *ANXA1* and *CTGF*. LDPE and HDPE exposure upregulated genes associated with neuronal development and differentiation, including *NTRK3, WNT2B, OTX2*, and *LMX1A*. Notably, LDPE exposure also increased *GRIN2A* and *ERBB4* expression and reduced *NRGN* expression, genes previously implicated in schizophrenia^19–21^.

**Figure 4:**
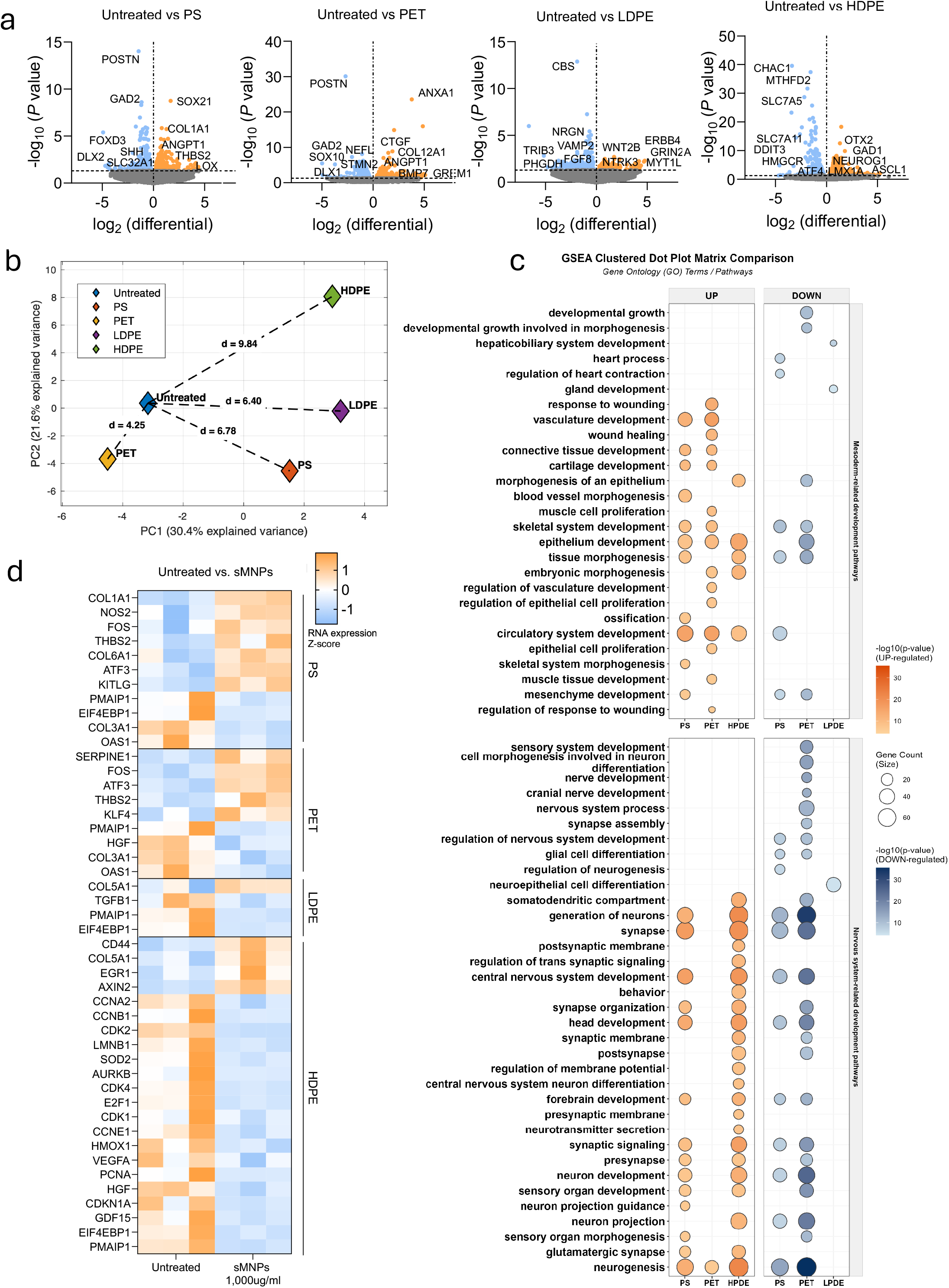
Polymer-specific sMNP exposure differentially alters neurodevelopmental and mesodermal transcriptional programs in developing COs. (**a**) Volcano plots depicting DEGs between untreated and 1,000 μg ml^-1^ PS, PET, LDPE, or HDPE-treated COs, exposed on day 6 of growth and collected 8 d.p.e. The x-axis shows log_2_ fold change, and the y-axis shows −log_10_(*P* value). Significantly upregulated genes are shown in orange, and significantly downregulated genes are shown in blue; nonsignificant genes are shown in gray. Representative genes associated with neurodevelopment, senescence, and extracellular matrix remodeling are indicated. Significant thresholds were set at p<0.05. (**b**) PCA of RNA-sequencing data in untreated and 1,000 μg ml^-1^ PS, PET, LDPE or HDPE treated COs, exposed on day 6 of growth and collected at 8 d.p.e. Each point represents an individual biological replicate, colored according to polymer exposure. PC1 (30.4% of explained variance) and PC2 (21.6% of explained variance) capture the major sources of transcriptomic variation. Diamonds represent group centroids, and dashed lines connect untreated and sMNP-treated centroids at each time point. Euclidean distances between centroids (untreated and sMNP-treated) quantify the separation between treatment groups. (**c**) GO enrichment analysis of significantly upregulated and downregulated DEGs in COs exposed to individual sMNP polymers (PS, PET, HDPE, or LDPE; 1,000 μg ml^-1^) compared with untreated controls at 30 d.p.e. Enriched GO terms associated with upregulated DEGs are shown in orange, and enriched GO terms associated with downregulated DEGs are shown in blue. Dot size corresponds to the number of genes associated with each GO term, and dot color represents enrichment significance (−log_10_ *P* value). (**d**) Heatmap of selected senescence-associated genes identified from the SenSig^13^ and SenMayo^14^ gene sets in untreated and PS, PET, LDPE, or HDPE-treated COs, exposed on day 6 of growth and collected 8 d.p.e. Gene expression is displayed as row-wise Z-scores of normalized RNA-sequencing data, with orange representing higher relative expression and blue representing lower relative expression.

PCA further demonstrated distinct transcriptional responses across polymers, with PC1 and PC2 accounting for 30.4% and 21.6% of the total variance, respectively (Fig. 4b). We observed separation between all polymer-treated groups and untreated controls, although the magnitude and direction of transcriptional divergence varied. HDPE exhibited the greatest centroid separation from untreated controls (d=9.84), followed by PS (d=6.78), LDPE (d=6.40), and PET (d=4.25). Importantly, we observed clear separation among each polymer in PCA space, indicating that individual sMNPs induce distinct rather than uniform transcriptional responses during early CO development. To identify biological processes affected by each polymer, we performed GO enrichment analysis on significantly upregulated and downregulated DEGs (Fig. 4c). Although each polymer exhibited a distinct enrichment profile, we identified several common biological themes. PS and PET predominantly enriched pathways associated with epithelial and connective tissue development and morphogenesis, whereas HDPE enriched pathways involved in nervous system development. PET also showed the broadest enrichment of downregulated neurodevelopmental processes, including neurogenesis, neuron differentiation, neuron projection development, synapse organization, gliogenesis, and central nervous system development. In comparison, LDPE showed few significantly enriched biological processes.

Given the polymer-specific transcriptional responses and enrichment of stress-associated biological processes, we next examined curated senescence-associated gene sets (SenSig^13^, SenMayo^14^) (Fig. 4d). Although the response differed among polymers, we identified several altered senescence-associated genes after acute exposure. HDPE exposure induced the greatest number of changes in senescence-associated gene expression, whereas PS, PET, and LDPE produced more modest but distinct transcriptional changes.

Overall, we found that individual sMNP polymers elicited distinct transcriptional and biological responses during early cortical development. Polymer-specific differences in gene expression, pathway enrichment, and senescence-associated signaling highlight how sMNP composition differentially affects cortical organoid development, with HDPE producing the most pronounced senescence-associated transcriptional changes and PET showing broad effects on neurodevelopmental pathways.

Having established that combined sMNP exposure disrupts neural progenitor maintenance, we next determined whether individual polymers contributed equally to these effects. In addition to differences in polymer composition, the particles displayed distinct morphologies: PS and HDPE consisting predominantly of spherical particles, whereas PET and LDPE were primarily irregular fragments (Supplementary Fig. 1a). After acute exposure, we observed polymer-specific effects on early CO development. PET and LDPE treatment significantly increased the amount of SOX2-positive neural progenitors relative to untreated organoids, whereas HDPE significantly reduced SOX2 expression (Supplementary Fig. 6a, b). PET and LDPE also significantly increased p21-positive cells (Supplementary Fig. 6a, b). ZO-1 expression within neuroepithelial structures decreased across COs exposed to all polymer types, indicating disruption of neuroepithelial cell-cell junctions that may alter intercellular signaling during cortical development, with PS producing the greatest reduction (Supplementary Fig. 6c, d). Although PAX6 expression was largely preserved following PS, PET, and LDPE exposure, we saw that HDPE exposure significantly reduced the proportion of PAX6-positive progenitors (Supplementary Fig. 6c, d), indicating that HDPE uniquely disrupts both neural stem cell maintenance and dorsal cortical progenitor populations during early cortical development.

Following persistent exposure, we observed more pronounced polymer-specific effects across markers of neural progenitor maintenance, neuronal maturation, and cellular senescence. PS, PET, and HDPE exposure led to unaltered SOX2-positive neural progenitors, whereas LDPE exposure significantly increased SOX2 expression at single-cell resolution (Supplementary Fig. 7a, b). In contrast to acute exposure, all polymer treatments reduced p21-positive cells after persistent exposure (Supplementary Fig. 7a, b). We also found polymer-specific effects on neuronal morphology and cytoskeletal organization with PS and HDPE exposure significantly reducing MAP2 expression, while PET and LDPE produced no significant changes (Supplementary Fig. 7c, d). HDPE exposure significantly reduced PAX6-positive progenitors, whereas PET exposure increased PAX6 expression (Supplementary Fig. 7e, f). Finally, p16 expression remained unchanged following PS, PET, and HDPE exposure but significantly increased following LDPE exposure (Supplementary Fig. 7e, f).

Overall, we found that individual sMNP polymers exert distinct, time-dependent effects on cortical development. Acute exposure produced divergent effects on neural progenitor maintenance and cell-cycle regulation, while persistent exposure more consistently reduced progenitor and neuronal markers, particularly with PS and HDPE. These findings highlight polymer composition as an important determinant of the cellular response to MNP exposure.

### eMNPs alter cortical neurogenic trajectories and promote Choroid Plexus-like states

Having established that sMNPs broadly disrupt neural progenitor maintenance and cortical development, we next examined whether environmentally derived micro- and nanoplastics (eMNPs) produced similar neurodevelopmental effects. We exposed COs to eMNPs on day 6 of differentiation and analyzed them after 30 days of exposure. We generated the eMNP preparation from plastic debris collected from Kamilo Beach, Hawaii, manually size-reduced the debris, washed it with soap and water followed by 100% ethanol, cryogenically milled it, and subjected it to 5 weeks of ultraviolet and ozone weathering to mimic environmental aging (Supplementary Fig. 8a). The weathered material was subsequently re-milled and size-fractionated to generate particles spanning multiple size ranges, including particles <20 µm with approximately 1% of the mass of eMNPs being submicron (Supplementary Fig. 8a). The resulting preparation contained a heterogeneous mixture of environmentally sourced polymers, with semicrystalline polyethylene and polypropylene representing the predominant polymer classes, alongside polyurethane, nylon 66, polymethyl methacrylate, polyvinyl chloride, PET, nylon 6, acrylonitrile butadiene styrene, styrene-butadiene rubber, polycarbonate, and PS (Supplementary Fig. 8b).

To further characterize the cellular response to eMNP exposure, we next examined how eMNP exposure altered the cellular composition of COs using single-nuclei sequencing (Fig. 5a-c and Supplementary Fig. 8c). eMNP exposure shifted the distribution of cortical developmental states, with a significant reduction in progenitor/newborn-neuron populations (∼46% to ∼44%; χ^2^(1) = 4.45, P = 0.0349) accompanied by a significant increase in immature neuronal populations (∼25.0% to ∼28%; χ^2^(1) = 29.04, P < 0.0001), similar to the effects of sMNPs (Fig. 5a, b and Supplementary Fig. 8c). This shift reflected reduced representation of several radial glial/progenitor states together with expansion of neurogenic-transition and immature neuronal populations, including an approximately 2-fold change in the neurogenic progenitor–newborn neuron transition population.

**Figure 5:**
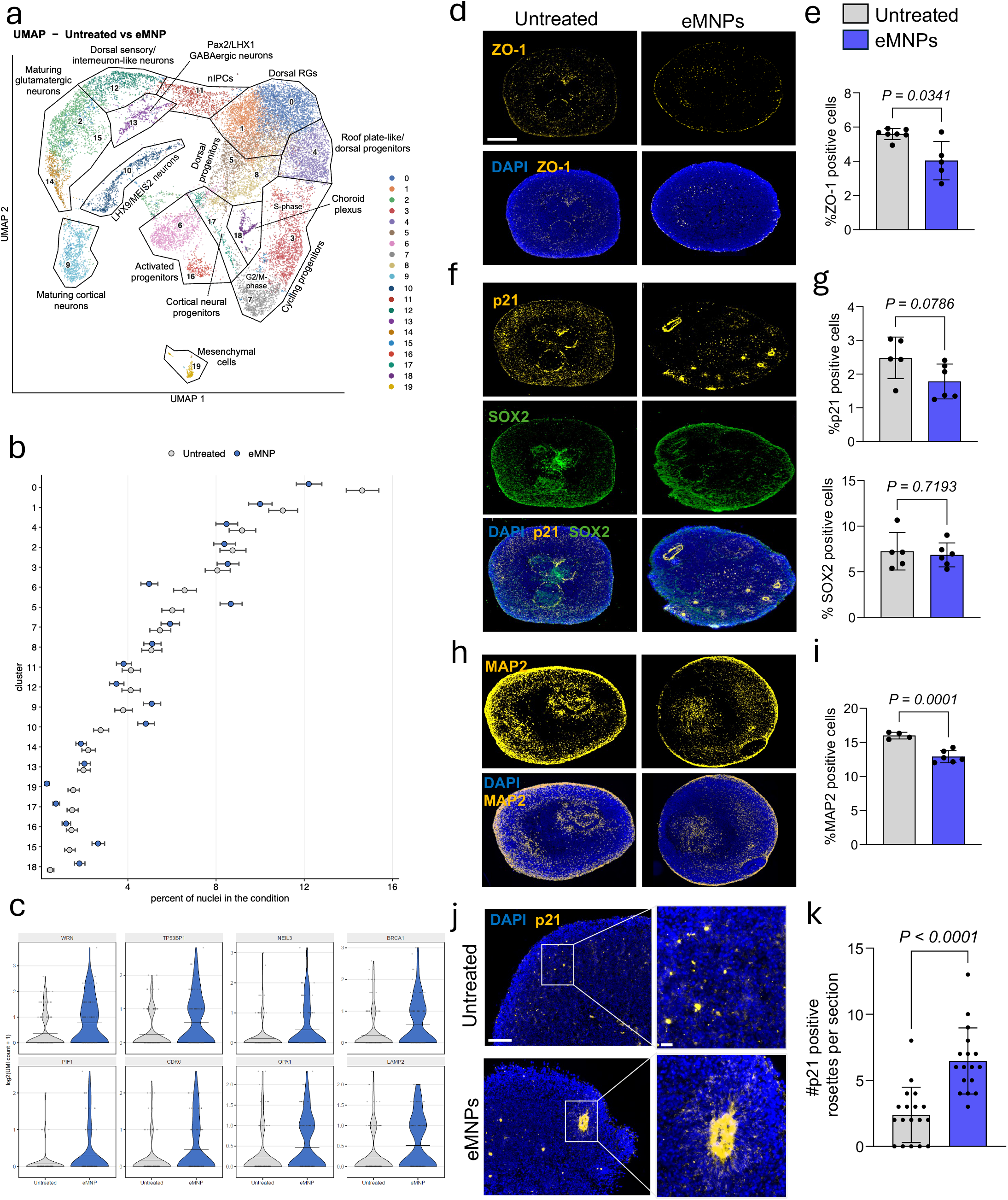
Environmental micro- and nanoplastics remodel cortical development and induce localized cellular stress. (**a**) UMAP visualization of single-nuclei RNA sequencing data from untreated and 1,000 μg ml^-1^ environmental micro- and nanoplastic (eMNP)-treated COs exposed on day 6 of growth and collected 30 d.p.e., showing the major transcriptionally defined cell populations identified across the dataset. Each point represents an individual nucleus, with colors indicating the corresponding cell cluster. Clusters were annotated based on established marker gene expression and include dorsal radial glia (RGs), neural intermediate progenitor cells (nIPCs), dorsal sensory/interneuron-like neurons, Pax2/LHX1 GABAergic neurons, maturing glutamatergic neurons, LHX6/MEIS2 neurons, cortical neural progenitors, activated progenitors, maturing cortical neurons, choroid plexus, roof plate-like/dorsal progenitors, cycling progenitors, and mesenchymal cells. (**b**) Relative composition of transcriptionally defined cell populations in untreated and 1,000 μg ml^-1^ eMNP-treated COs following 30 d.p.e. Data are presented as the proportion of nuclei assigned to each cell cluster within each treatment condition. Comparisons are based on relative proportions rather than raw cell counts because the total number of captured nuclei differed between conditions. Because each treatment condition was represented by a single 10x capture, differences in cell-type composition are presented as descriptive, hypothesis-generating observations and were not statistically tested for treatment effects. (**c**) Violin plots showing expression of representative differentially expressed genes in cluster 17, corresponding to cortical neural progenitors, from untreated and 1000 μg ml^-1^ eMNP-treated COs exposed on day 6 of growth and collected 30 d.p.e. eMNP exposure was associated with increased expression of genes involved in DNA damage and genome maintenance (*WRN, TP53BP1, NEIL3, BRCA1, PIF1*), cell-cycle regulation (*CDK6*), and mitochondrial and lysosomal homeostasis (*OPA1, LAMP2*). Gene expression is shown as log_2_-transformed UMI counts + 1. Each violin represents the distribution of expression across cells within the indicated condition. **d,f,h,** Representative immunofluorescence images of untreated and 1,000 μg ml^-1^ eMNP-treated COs, exposed on day 6 of growth and collected 30 d.p.e. Sections were stained for ZO-1 (yellow) (**d**); p21 (yellow) and SOX2 (green) (**f**); and MAP2 (yellow) (**h**). Nuclei were counterstained with DAPI (blue). Merged images are shown for each staining panel. Scale bars = 400 μm. **e,g,i,** Quantification of ZO-1-positive (**e**); p21-positive and SOX2-positive (**g**); and MAP2-positive (**i**) cells in untreated and eMNP-treated COs, exposed on day 6 of growth and collected 30 d.p.e. Data are presented as mean ± SD, with each data point representing an individual organoid. Statistical significance was determined using a two-tailed Welch’s *t*-test. Exact *P* values are shown. n = 5-7 organoids per treatment group from a single organoid differentiation. (**i**) Representative immunofluorescence image of a neural rosette in untreated and COs exposed to eMNPs on day 6 of growth and collected 30 d.p.e., stained for p21 (yellow) and DAPI (blue). The circled region highlights a representative p21 positive rosette, with the corresponding higher-magnification image shown at right. Scale bars= 100 μm for lower magnification and 20 μm for higher magnification. (**j**) Quantification of the number of p21-positive rosettes per organoid section in untreated and eMNP-treated COs, exposed on day 6 of growth and collected 30 d.p.e. Each data point represents an individual organoid section. Data are presented as mean ± SEM. Statistical significance was determined using an unpaired two-tailed Student’s *t*-test. Exact *P* values are shown.

Most notably, eMNP exposure was associated with a nearly 4-fold enrichment of the choroid plexus/dorsal midline-like population (Fig. 5b and Supplementary Fig. 8c). This population exhibited marked induction of *TTR* following eMNP exposure (log2FC = 5.14, FDR = 0.011), further supporting a shift toward a choroid plexus-like transcriptional state. We also identified pronounced transcriptional remodelling within cortical neural progenitors (cluster 17), characterized by increased expression of genes involved in DNA damage and genome maintenance (*WRN, TP53BP1, NEIL3, BRCA1, PIF1*), cell-cycle regulation (*CDK6*), and mitochondrial and lysosomal homeostasis (*OPA1, LAMP2*) (Fig. 6c). These changes indicate that eMNP exposure is associated with increased replication/genotoxic stress and senescence within cortical neural progenitors. Together, these findings indicate that eMNP exposure alters cortical neurogenic trajectories while preferentially promoting a choroid plexus/dorsal midline-like cellular state.

**Figure 6:**
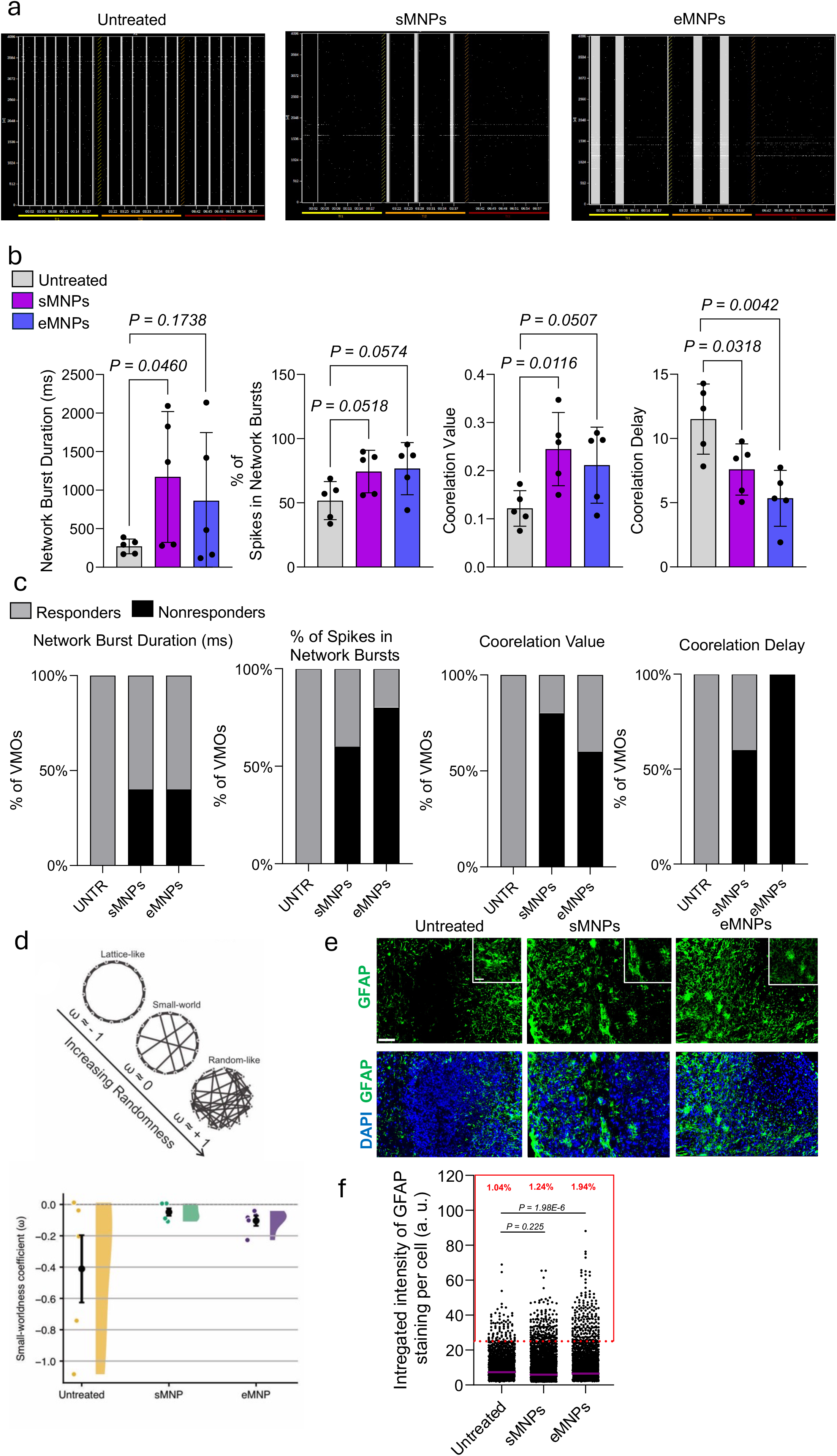
MNP exposure alters neuronal network functions in mature ventral midbrain organoids. (**a**) Representative multi-electrode array (MEA) raster plots from untreated, sMNP, or eMNP treated ventral midbrain organoids (VMOs), exposed on day 90 of growth and collected 30 d.p.e. The x-axis represents recording time (min), and the y-axis represents individual electrodes in the MEA array. Each vertical line represents a detected neuronal spike across active electrodes during the recording period. Raster plots illustrate differences in network burst synchronization between treatment groups. (**b**) Quantification of MEA network activity, including network burst duration (ms), percentage of spikes within network bursts, correlation value, and correlation delay of untreated, sMNP or eMNP treated VMOs. Bars represent the mean ± SEM, with each point representing an individual organoid. Statistical significance was determined using unpaired Student’s *t*-tests comparing each treatment group with untreated controls. (**c**) Stacked bar graphs showing the percentage of responder and nonresponder untreated, sMNP or eMNP-treated VMOs for each MEA network activity metric. Organoids were classified as responders or nonresponders based on predefined threshold values for each individual parameter. (**d**) The small-worldness coefficient (ω) was used to compare network topology with lattice-like and random reference networks. Values approaching −1 indicate lattice-like organization, values near 0 indicate small-world organization, and values approaching +1 indicate random organization. sMNP-exposed organoids exhibited ω values clustered near 0, consistent with small-world-like topology, whereas eMNP-exposed organoids showed a narrower distribution with a slight shift toward lattice-like organization. Untreated organoids exhibited greater variability in ω values. (**e**) Representative higher-magnification immunofluorescence images of untreated, 1,000 μg ml^-1^ sMNP-, or eMNP-treated VMOs, exposed on day 90 of growth and collected 30 d.p.e. Sections were stained for GFAP (green), and nuclei were counterstained with DAPI (blue). Merged images are shown for each treatment group. Higher magnification panels highlight astrocyte formation. Scale bars = 50 μm for lower resolution and 20 μm for higher resolution. (**f**) GFAP immunofluorescence intensity quantified at the single-cell level in untreated, sMNP-, and eMNP-treated cortical organoids. GFAP-high cells were defined as cells with integrated GFAP staining intensity >25 arbitrary units, corresponding to the 99th percentile of the untreated population (red dashed line). Each point represents an individual cell. The percentage of GFAP-high cells was calculated for each treatment group. Statistical significance was determined using a chi-square test comparing the frequency of GFAP-high and non-GFAP-high cells across treatment groups, followed by pairwise chi-square comparisons. Exact *P* values are shown.

We next examined whether these cellular changes were accompanied by alterations in cortical tissue organization and cellular identity at the protein level. We found that eMNP treatment significantly reduced the proportion of ZO-1-positive cells (Fig. 5d, e) and did not significantly alter the proportion of p21 and SOX2-positive cells (Fig. 5f, g). In contrast, eMNP exposure significantly reduced MAP2-positive neurons (Fig. 5h, i), consistent with impaired neuronal maturation/maintenance. Although global p21 expression was not increased, we examined high-resolution analysis of p21, revealing a localized redistribution of expression within the tissue. Representative immunofluorescence imaging showed p21-positive neural rosettes in eMNP-treated organoids, with p21 signal concentrated within discrete neuroepithelial structures (Fig. 5j). Quantification confirmed a significant increase in the number of p21-positive neural rosettes per organoid section following eMNP exposure (Fig. 5k), indicating that p21-mediated cell cycle arrest is spatially concentrated within neuroepithelial rosettes, rather than broadly distributed throughout the organoid.

Overall, we found that prolonged eMNP exposure alters cortical tissue organization and neuronal maturation while changing the relative abundance of specific progenitor and neuronal populations, including increased representation of newborn-neuron transition and several immature neuronal populations. eMNP exposure also produced localized p21 activation within neuroepithelial rosettes. These findings indicate that environmentally derived MNPs induce spatially restricted cellular stress and neurodevelopmental alterations while preferentially expanding a choroid plexus/dorsal midline-like transcriptional state.

### MNP exposure alters neuronal network function in mature brain organoids

To determine whether the effects of micro- and nanoplastics (MNPs) extend beyond cortical development, we exposed mature COs to sMNPs or eMNPs on day 90 of differentiation and analyzed them after 30 days of prolonged exposure. Neither sMNP nor eMNP altered ZO-1 expression (Supplementary Fig. 9a, b). In contrast, sMNP exposure significantly increased TTR-positive cells and NeuN-positive mature neurons, whereas eMNP exposure did not significantly alter either marker (Supplementary Fig. 9a-d). Despite the increase in p21-positive neural rosettes observed with eMNP exposure (Fig. 5j, k), both sMNP and eMNP exposure significantly reduced the percent of p21-positive cells, suggesting redistribution of p21-positive cells rather than a global increase in p21 expression (Supplementary Fig. 9c, d). Both MNPs also significantly reduced SOX2-positive cells, while sMNP exposure significantly increased MAP2-positive neurons and eMNP exposure produced no significant change (Supplementary Fig. 9e, f). Finally, sMNP exposure resulted in a trend towards reduced GFAP-positive astrocytes, whereas eMNP-treated organoids were comparable to untreated controls (Supplementary Fig. 9g, h). Neither MNP exposure significantly altered p16 expression, revealing no evident signs of senescence (Supplementary Fig. 9g, h).

We next examined a distinct brain region using mature ventral midbrain organoids (VMOs). We exposed VMOs to sMNPs or eMNPs on day 90 of differentiation and analyzed them following 30 days of prolonged exposure. Neither MNP exposure produced statistically significant changes in TH-positive dopaminergic neurons, LMX1A-positive ventral midbrain progenitors, p21-positive, TTR-positive, or SOX2-positive cells (Supplementary Fig. 10a-f), indicating that prolonged MNP exposure produced only modest effects on cellular composition.

To further quantify changes in GFAP expression at the single-cell level, we identified GFAP-positive cells and quantified their integrated staining intensity (Fig. 6e, f). We classified cells with integrated GFAP staining intensity >25 arbitrary units as GFAP-high, corresponding to the 99th percentile of the untreated population (Fig. 6f). GFAP-high cells comprised 1.04% of cells in untreated VMOs, compared with 1.24% following sMNP exposure and 1.94% following eMNP exposure. eMNP exposure significantly increased the proportion of GFAP-high cells, indicating that eMNPs promote a localized increase in cells with elevated GFAP expression, and may enhance astroglial responses to MNP exposure.

To determine whether MNP exposure altered neuronal function despite the limited changes in VMO cellular identity, we assessed spontaneous neuronal activity using multi-electrode array (MEA) recordings (Fig. 6a-d and Supplementary Fig. 11a). Representative raster plots showed differences in the temporal organization of spontaneous neuronal activity following MNP exposure, with sMNP- and eMNP-treated VMOs exhibiting more pronounced periods of coordinated network activity compared with untreated organoids (Fig. 6a).

We quantified network activity and saw that sMNP exposure significantly increased network burst duration compared with untreated organoids, indicating more sustained periods of synchronized network activity, whereas eMNP exposure produced a similar but non-significant trend (Fig. 6b). The percentage of spikes occurring within network bursts also increased following both sMNP and eMNP exposure, although neither reached statistical significance (Fig. 6b). sMNP exposure significantly increased network correlation values, with a similar trend following eMNP exposure, suggesting enhanced synchronization of neuronal activity (Fig. 6b). In contrast, both sMNP and eMNP exposure significantly reduced correlation delay, consistent with a more rapid propagation of synchronized network activity across the neuronal network (Fig. 6b). Analysis of individual organoids as responders or nonresponders further showed that a greater proportion of MNP-exposed organoids met responder criteria for network burst duration, percentage of spikes in network bursts, correlation value, and correlation delay compared with untreated organoids (Fig. 6c), supporting a reproducible shift toward altered network dynamics following both sMNP and eMNP exposure.

We further evaluated network organization using the small-worldness coefficient, which describes the balance between locally clustered and globally interconnected network architecture (Fig. 6d). MNP exposure altered network architecture, with treated organoids exhibiting small-worldness values closer to zero than untreated controls (Fig. 6d). Representative network connectivity maps further illustrate differences in the organization across functional electrodes following MNP exposure (Supplementary Fig. 11a).

Overall, prolonged MNP exposure produced relatively modest changes in cellular composition but more pronounced alterations in neuronal network function. sMNPs and eMNPs altered neural progenitor and astroglial markers in a context-dependent manner, while MEA recordings revealed changes in network bursting, synchronization, propagation, and organization. These findings suggest that functional alterations in neuronal networks may represent a more sensitive indicator of MNP-induced neurotoxicity than changes in cellular identity alone.

## Discussion

MNPs are increasingly detected in human tissues, including the brain, but their functional consequences from acute and chronic exposure remain unknown^2^. Here, we establish MNP vulnerability as a multifaceted perturbation of human neurobiology, simultaneously altering developmental trajectories, cellular senescence and neuronal network function, suggesting that the effects of MNP exposure may extend well beyond transient cellular stress.

Human brain development requires the precise specification of embryonic ectoderm into neuroectoderm, followed by the coordinated expansion and differentiation of neural progenitors into increasingly specialized neuronal and glial populations^22–24^. Disruption of these early developmental programs can have consequences that persist long after the initial insult^25^, and altered neurodevelopmental trajectories (including biases towards Choroid Plexus formation) have been implicated in disorders ranging from autism spectrum disorder and schizophrenia to Alzheimer’s disease^26–28^. Here, we find that MNP exposure profoundly disrupts this developmental landscape. Exposure suppressed neuroectodermal and neuronal programs while increasing mesoderm-associated developmental signatures, indicating a shift away from normal neural specification. Single-nucleus profiling further resolved this disruption at the cellular level, revealing altered representation of cortical progenitor and neuronal states and, particularly following eMNP exposure, a marked enrichment of a TTR-high choroid plexus/dorsal midline-like population. This unexpected redirection of developmental identity suggests that MNP exposure interferes with lineage decisions at fundamental stages of human neurodevelopment.

This developmental disruption was accompanied by a progressive senescence-associated response. Acute MNP exposure activated stress, inflammatory and cell-cycle pathways, whereas prolonged exposure produced stronger enrichment of senescence-associated programs together with increased p21 expression and depletion of neural progenitor populations. Single-nucleus analyses further revealed that these responses were cell-state specific, with sMNP-exposed maturing glutamatergic neurons displaying inflammatory, oxidative and DNA damage-associated transcriptional programs, while eMNP-exposed cortical neural progenitors exhibited coordinated activation of genome-maintenance and mitochondrial/lysosomal homeostasis pathways associated with cellular stress and aging. These findings suggest that MNP exposure may couple developmental disruption to a senescence-associated state with consequences extending beyond the period of exposure. Premature senescence during development could restrict progenitor expansion and differentiation while establishing a pro-inflammatory microenvironment capable of propagating senescence or dysfunction to neighboring cells^29–31^. Such persistent alterations may compromise normal brain maturation, raising the possibility that MNP exposure during critical developmental windows could establish cellular vulnerabilities that manifest as neurological dysfunction later in life^32^.

Importantly, the consequences of MNP exposure extended beyond early developing tissue. In mature midbrain organoids, MNP exposure altered neuronal firing synchronization, burst dynamics and the propagation of network activity despite relatively modest changes in cellular composition. Rather than reflecting enhanced neuronal function, increased synchronization represents a deviation from normal patterns of coordinated activity and altered network function^33^. Thus, MNP-induced cellular and developmental perturbations may ultimately converge on network-level dysfunction, providing a potential functional pathway through which early and prolonged exposure could alter brain function.

Limitations to the study should be considered in interpreting these findings. While we tested an array of polymers, including environmentally sourced MNPs, the exposures were dose limited. Biologically speaking, there is still considerable uncertainty regarding internal human doses. Nihart et al reported total plastics concentrations of approximately 5,000 µg g^-1^ in the frontal cortex of decedents from New Mexico, USA^2^. More recently, Li et al reported much lower concentrations (∼130 µg g^-1^) in different regions of the brain adjacent to tumors^3^, although these samples were from participants from China, where the historic usage of plastics is a small fraction (∼3-10%) of the usage in the USA, even though they are comparable in modern times^34,35^. In terms of the present study, 1,000 µg ml^-1^ would sit in between these estimates from human samples. Notably, in the Nihart study there appeared to be focused regions or “hotspots” of likely plastics in individual cells, thus there may be a wide range of concentrations between regions and cell types. Lastly, the use of the environmental MNP model does not perfectly reflect the proportions and morphologies of true environmental nanoplastics. However, this material does better represent the weathering that occurs due to sunlight, heat, and other factors, in addition to representing mixed material that has actually been created, marketed, used, and disposed of, as compared to pristine synthetic microspheres. By using the varied models in complement, we can reveal common outcomes that are likely to be of greatest concern. Collectively, our findings highlight MNPs as an increasingly important environmental concern with the potential to affect human brain health across early and late developmental and mature neural states. The ability of MNPs to disrupt neural lineage specification, promote a persistent senescence-associated phenotype and alter neuronal network activity, even in the absence of overt cytotoxicity, underscores the need to recognize MNP exposure as a potential contributor to neurological risk. Although the precise mechanisms linking MNP exposure to these phenotypes remain to be defined, our findings provide a foundation for investigating how particle properties, exposure timing and cellular responses interact to drive neural dysfunction. Continued investigation of these mechanisms, including studies of long-term exposure and individual susceptibility will be essential to determine the broader neurological consequences of this increasingly pervasive environmental contaminant.

## Materials and methods

### Human iPSC culture

Healthy control Human Induced Pluripotent Stem Cell (hiPSC) line SCTiOO3-A (STEMCELL Technologies, cat. #200-0511) were generated from PBMCs of one female patient. All human pluripotent stem cell lines were routinely tested and confirmed negative for mycoplasma (MycoAlert, Lonza). hiPSCs were maintained under feeder-free conditions on Matrigel (Thermo Fisher cat. # 8774552)-coated six-well tissue culture plates in mTeSR™ Plus medium (STEMCELL Technologies, cat. # 100-0276) at 37°C with 5% CO₂. Tissue culture plates were coated with Matrigel diluted in DMEM/F-12 (Thermo Fisher, cat. # 11330057) according to the manufacturer’s instructions and incubated for a minimum of 30 min at 37°C prior to cell seeding. Cryopreserved hiPSCs were rapidly thawed and transferred to mTeSR™ Plus medium supplemented with ROCK inhibitor (Thermo Fisher, cat. # 501031738). Cells were centrifuged at 150 × *g* for 5 min, resuspended in fresh mTeSR™ Plus containing ROCK inhibitor, and seeded dropwise onto Matrigel-coated six-well plates. Cells were maintained at 37°C and 5% CO₂, and the culture medium was replaced daily with fresh mTeSR™ Plus medium. ROCK inhibitor was included only during the initial plating following thawing.

### Cortical organoid generation

Human induced pluripotent stem cells (hiPSCs) derived from a female donor (SCTiOO3-A) were differentiated into cortical organoids using a modified feeder-free protocol based on the method described in the Pasca et al protocol^36^. Briefly, hiPSCs were dissociated into single cells using Accutase (Sigma-Aldrich, cat. # A6964) and seeded into ultra-low attachment 96-well plates (Thermo Scientific, cat. # 7201680) at a density of 10,000 cells per well in mTeSR™ Plus medium (STEMCELL Technologies, cat. # 100-0276) supplemented with Culture-CEPT (Thermo Fisher, cat. # A56800) to generate embryoid bodies. Plates were centrifuged at 100 × *g* for 3 min and maintained at 37°C with 5% CO₂. On day 0, embryoid bodies were transferred to anti-adherent 100-mm culture dishes containing Essential 6 (E6) medium (Thermo Fisher, cat. #A1516401) supplemented with 2.5 μM dorsomorphin (Sigma-Aldrich, Cat. No. P5499-5MG) and 10 μM SB431542 (Sigma-Aldrich, cat. #S4317-5MG). Organoids were maintained on an orbital shaker at 85 rpm, and the medium was replaced daily through day 5. Beginning on day 6, organoids were transferred to ultra-low attachment six-well plates and cultured in neural medium consisting of Neurobasal™ medium (Thermo Fisher, cat. #21103049) supplemented with B-27™ Supplement Minus Vitamin A (Thermo Fisher cat. #12587001), GlutaMAX™ Supplement (Thermo Fisher, cat. #35050061), and 1% penicillin-streptomycin (Thermo Fisher, cat. #15140122). Neural medium was additionally supplemented with 20 ng ml^-1^ epidermal growth factor (EGF; Thermo Fisher, cat. #AF10015100) and 20 ng ml^-1^ fibroblast growth factor 2 (FGF2; Thermo Fisher, cat. #10018B50UG) from days 6–24, with daily media changes from days 6–17 and every other day from days 18–24. From days 25–42, organoids were maintained in neural medium supplemented with 20 ng ml^-1^ brain-derived neurotrophic factor (BDNF; Thermo Fisher, cat. #4500250UG) and 20 ng ml^-1^ neurotrophin-3 (NT-3; Thermo Fisher, cat. # 34500350UG), with media exchanged every other day. Beginning on day 43, organoids were cultured in neural medium without exogenous growth factors, and media were replaced every four days or as needed until experimental endpoints.

### Ventral midbrain organoid generation

Ventral midbrain organoids were generated from human induced pluripotent stem cells (STEMCELL Technologies, cat. #200-0511), using the Reumann et al protocol^37^. For organoid generation, 9,000 cells were seeded in 150 µl neural induction medium supplemented with 200 ng ml^-1^ Noggin (Thermo Fisher, cat. # 6057NG100), 10 µM SB431542 (Sigma, cat. no. 616464), 1 µM CHIR99021 (Sigma, cat. no. 361571), and ROCK inhibitor (Thermo Fisher cat. # 501031738) in ultra-low attachment U-shaped 96-well plates. On day 2, 100 µl of medium was removed and replaced with 150 µl of fresh medium of the same composition. On day 4, medium was exchanged and supplemented with 200 ng ml^-1^ Noggin, 10 µM SB431542, 1 µM CHIR99021, 300 nM SAG (Sigma, cat. #566661), and 100 ng ml^-1^ FGF-8 (Thermo Fisher, cat. # 5027FF025). On day 6, medium was replaced with neural induction medium containing 300 µM SAG and 100 ng ml^-1^ FGF-8. On day 8, up to 30 embryoid bodies were transferred to 10 cm plates coated with anti-adherence rinsing solution (STEMCELL Technologies, cat. #07010) and maintained in 12 ml Improved-A medium supplemented with 2% liquid Matrigel (Thermo Fisher cat. # 8774552) prepared in pre-chilled medium and used within 1 h, with a final exposure to 300 µM SAG and 100 ng ml^-1^ FGF-8 at the first feeding after transfer. Improved-A medium consisted of a 1:1 mixture of DMEM/F12 (Thermo Fisher, cat. # 11330057) and Neurobasal (Thermo Fisher, cat. no. 21103049) supplemented with 0.5% N2, 2% B27−A (Thermo Fisher cat. #12587001), insulin (1:4,000; Sigma-Aldrich cat. #I9278), 1% GlutaMAX (Thermo Fisher, cat. #35050061), 0.5% MEM-NEAA (Thermo Fisher, cat. no. 11140050), and 1% Antibiotic-Antimycotic (Thermo Fisher, cat. no. 15240062). Following day 8, organoid medium was exchanged every 3 days, with a transition to Improved+A medium around day 16. Improved+A medium consisted of a 1:1 mixture of DMEM/F12 and Neurobasal (Thermo Fisher, cat. #21103049) supplemented with 0.5% N2 (Thermo Fisher, cat. #17502001), 2% B27+A (Thermo Fisher, cat. #17504001), insulin (1:4,000; Sigma-Aldrich cat. #I9278), 1% GlutaMAX, 0.5% MEM-NEAA, 1% Antibiotic-Antimycotic (Thermo Fisher, cat. #15240062), 1% vitamin C solution prepared from a 40 mM stock in DMEM/F12 (Vitamin C, Sigma-Aldrich, cat. # A4544), and 1 g l−1 sodium bicarbonate (Sigma-Aldrich, cat. #S5761). Organoids were maintained under static conditions until day 20, after which they were transferred to orbital shakers at a reduced speed of 85 rpm.

### Synthetic micro- and nanoplastic oxidation

Polyethylene terephthalate (PET), high density polyethylene (HDPE), and low-density polyethylene (LDPE) powders, and polystyrene (PS) microspheres were purchased from Nanochemazone (NCZ-PA-138/24, NCZ-PE-120/0725B, NCZ-AE-120/0725C, NCZ-NP-652/24). The plastics were described by manufacturers as 1-5 μm in size. Plastics were treated with low-intensity UVA (365nm) irradiation for 72 h with a Spectroline E-Series UV-A Lamp (Sigma, Z169595), which has a typical peak intensity of 300 μW/cm2. Plastics were spread in thin layer in petri dish and irradiated for 72 h in a closed chamber with stirring every 8 hr. The dish was located 8 cm from the light source. The UV-treated plastic samples were washed with ethanol, dried overnight, and collected for future experiments. UV weathering was found to improve the dispersibility in aqueous media of microplastic particles. Microplastic size distribution was assessed by dynamic light scattering (DLS). Microplastics were dispersed in MilliQ water with 0.05% Tween 20 to increase dispersibility, then sonicated for 10 minutes at 50% amplitude in a Qsonica Q500 Sonicator with VWR circulating water bath. Microplastic suspensions were measured for zeta potential using a Malvern ZetaSizer Ultra instrument (Malvern Panalytical, Westborough, MA, USA). Plastic polymers were confirmed with Attenuated Total Reflectance Fourier-Transform Infrared Spectroscopy (ATR-FTIR). A small sample of microplastic was placed on the built-in diamond ATR crystal on a Thermo Scientific Nicolet iS50 FTIR spectrometer. FTIR spectra were collected from 400-4000nm using 64 scans for sample and background measurements.

### Environmental micro- and nanoplastic particle preparation

Plastic debris was collected from Kamilo Beach on the southern coast of Hawai’i Island. Kamilo Beach is located within the North Pacific Subtropical Gyre, a convergence zone influenced by the Kuroshio, North Pacific, California, and North Equatorial currents, resulting in substantial accumulation of marine plastic debris. Approximately 50 lbs. of plastic material was collected and transported by the Campen Lab for processing. Collected plastic pieces were manually reduced in size to dimensions suitable for cryogenic milling. Fragments were washed with soap and water for 24 hours using a stir plate to remove adhered sediment and environmental residues, followed by immersion in 100% ethanol for another 24 hours to further reduce surface contamination. Cleaned plastic fragments were transferred into a 50 ml cryomill jar and processed under cryogenic conditions using a Retsch cryomill (Retsch, Haan, Germany) for a total of three cycles each consisting of 2 min of cooling at 5Hz and then 30 s of grinding at 30 Hz to generate micro- and nanoplastic particles. The cryomilled plastics then underwent accelerated weathering through ultraviolet and ozone exposure over a 5-week total time period, with one-week intervals of ultraviolet only and ozone only cycles to simulate additional environmental oxidative degradation. Following the 5 weeks of aging, plastics were reprocessed by cryogenic milling using a 50 ml cryomill jar and subjected to a dry sieving step to separate particles into three size fractions: 500 µm, 250 µm, and 125 µm Particles collected within the 125-µm fraction were selected for further refinement. This fraction underwent wet sieving using 100% ethanol to isolate particle size fractions 63 µm and the final size for utilization of less than 20 µm. Particles isolated to less than 20 µm fractions were collected and allowed to dry following ethanol extraction before being prepared for incorporation into experimental designs for in vivo and in vitro study models. Dynamic light scattering suggested that approximately 1% of the mass of eMNPs were in submicron range.

### MNP exposure

CO and VMOs were exposed to synthetic micro- and nanoplastic (sMNPs) or environmental micro- and nanoplastic (eMNP) treatments through supplementation of the culture medium with particle suspensions. Untreated control (UNTR) organoids received an equivalent volume of phosphate-buffered saline (PBS, Sigma-Aldrich, cat # D8537). Polystyrene (PS), polyethylene terephthalate (PET), low-density polyethylene (LDPE), high-density polyethylene (HDPE), and environmental ocean micro- and nanoplastics (eMNPs) were obtained as dry powders. Particle suspensions were prepared in sterile PBS and mixed on a rotary shaker at 4°C for 24 h to facilitate particle dispersion. Suspensions were subsequently stored at 4°C and vortexed immediately prior to use. Treatment groups consisted of individual exposures to PS, PET, LDPE, HDPE, or eMNPs at a final concentration of 1,000 μg ml^-1^. Additional groups were exposed to mixed-plastic suspensions at final concentrations of 10 μg ml^-1^ or 1,000 μg ml^-1^. The 1,000 μg ml^-1^ mixed-plastic treatment contained equal concentrations of each polymer type, including 250 μg ml^-1^ PS, 250 μg ml^-1^ PET, 250 μg ml^-1^ LDPE, and 250 μg ml^-1^ HDPE, resulting in a total MNP concentration of 1,000 μg ml^-1^. Organoids were maintained on an orbital shaker at 85 rpm throughout the exposure period to promote continuous contact between the organoids and suspended particles and to minimize particle settling. MNP exposures were administered every 7 days by adding the appropriate volume of particle suspension directly to the culture medium to achieve the desired final concentration.

### Multi-electrode array recordings and data analysis

Electrophysiological activity of VMOs was assessed at day 120 of differentiation using a BioCAM DupleX high-density multielectrode array (MEA) system (3Brain AG) equipped with a 64 × 64 electrode array, each electrode being a micropillar of 90 μm in height. Prior to each recording, the MEA chip was rinsed twice with PBS. Individual organoids were transferred from culture plates using wide-bore pipette tips and carefully positioned over the center of the recording array. A tissue anchor (3Brain AG) was placed over each organoid to maintain stable contact with the electrode array, and the recording chamber was filled with their respective culture medium. Recordings were performed at 37°C using BrainWave 6 software (3Brain AG). To minimize experimental variability, the interval between removal of each organoid from the incubator and the start of electrophysiological recording was standardized to exactly 7 min for all samples. All recordings were performed using identical acquisition settings. Spontaneous neuronal activity was recorded for 7 min for each organoid. Raw electrophysiological recordings were exported for downstream spike detection, burst analysis, and quantitative electrophysiological analyses. Spikes were detected using the peak-time standard deviation (PTSD) algorithm with a standard deviation factor of 8, a peak lifetime period of 2 ms, and a refractory period of 1 ms; spike assignment was performed automatically. Spike sorting was performed using principal component analysis (PCA) with three features, followed by K-means clustering with Gap Statistic optimization. A minimum of two spikes per cluster and a maximum of three clusters were permitted. Outliers were discarded using an outlier threshold of 2, and duplicate units were not removed. Spike bursts were identified using a maximum inter-spike interval of 100 ms and a minimum of five spikes per burst. Network bursts were detected using the recruitment method with a valid unit threshold of 0.1 spikes s⁻¹, a recruited valid unit threshold of 10%, a recruited spike threshold of 50, and a bin size of 50 ms. Data was analyzed by taking the mean number of events occurring over the entire recording period and graphics made using GraphPad Prism (version 11; GraphPad Software). Functional-network analysis was performed in Python using an approach adapted from MEA-NAP^9^. Artifact-cleaned spike trains exported from BrainWave were used to calculate pairwise functional connectivity with the spike-time tiling coefficient (STTC). A 25-ms coincidence window was selected as the primary analysis setting, with 10- and 50-ms windows examined as sensitivity analyses. Connections were retained when their observed STTC exceeded the 95th percentile of 180 independently time-shifted surrogate comparisons. The resulting networks were used to calculate network density, the mean of the strongest 10% of edge weights, community structure using consensus Louvain module detection, the mean of the highest 10% of participation coefficients, and the small-worldness coefficient omega. Each recording represented one independent organoid. The analysis followed the metric definitions and general framework described in MEA-NAP but was implemented independently in Python to accommodate the BrainWave-derived data and HD-MEA format.

### Tissue processing and cryosectioning

Organoids were washed once with PBS and fixed in 4% paraformaldehyde (PFA, Fujifilm Biosciences cat. # 16320145) for 1 h at 4°C. Following fixation, organoids were cryoprotected in 30% (w/v) sucrose (Sigma-Aldrich, cat. #S9378) in PBS for up to 5 h at 4°C. Organoids were then transferred to cryomolds containing a 40:60 mixture of optimal cutting temperature (O.C.T. Tissue-Tek cat. #4583) compound and 30% sucrose solution and frozen on dry ice. Embedded organoids were stored at −80°C until cryosectioning. Frozen organoids were cryosectioned on a Leica CM1850 at a thickness of 12 μm using a cryostat and mounted onto Superfrost™ Plus microscope slides (Thermo Fisher, cat. # 12-550-15). Tissue sections were stored at −20°C until immunofluorescence staining.

### Immunofluorescence staining

Immunofluorescence staining was performed at room temperature in a light-protected humidified chamber. A hydrophobic barrier pen was used to outline each tissue section and allowed to dry for 10 min. Sections were permeabilized with 0.2% Triton X-100 (Sigma Aldrich cat. # X100) in PBS for 10 min, followed by blocking with 1× PBG blocking solution (0.5% Bovine Serum Albumin (BSA) (Avantor cat. #97061-420) and 0.2% Gelatin from cold water fish skin (Sigma-Aldrich cat. #G7765-1L) in PBS) for 1 h. Primary antibodies (Table S1) were diluted in 1× PBG at optimized concentrations and incubated with tissue sections overnight. The following day, sections were washed four times with PBS for 7 min each before incubation with species-specific fluorescent secondary antibodies (Table S2) diluted 1:500 in 1× PBG for 45 min. Antibody combinations used for each immunofluorescence panel are listed in Table S3. Sections were then washed four times with PBS for 7 min each. Nuclei were counterstained with 0.2 μg ml^-1^ DAPI (Sigma Aldrich, cat. # D9564) in PBS for 2 min, followed by a final 7 min wash with PBS. Coverslips were mounted using ProLong™ Glass Antifade Mountant (Thermo Fisher, cat. # P36984) and allowed to cure overnight at room temperature in the dark before storage at 4°C until imaging.

### Antibodies

The following were used: anti-ZO-1 (Thermo Fisher, 1:200, cat. # 339100); anti-SOX2 (Cell Signaling Technology, 1:400, cat. # 4900S); anti-p21 (Thermo Fisher, 1:500, cat. # AF1047); anti-PAX6 (Thermo Fisher, 1:500, cat. # AF8150); anti-p16 (Cell Signaling Technology, 1:300, cat. # 80772S); anti-NeuN (Sigma-Aldrich, 1:1,000, cat. # MAB377); anti-MAP2 (Cell Signaling Technology, 1:200, cat. # 4542S); anti-TTR (Proteintech, 1:500, cat. # 11891-1-AP); anti-GFAP (Thermo Fisher, 1:1,000, cat. # 130300); anti-TH (Abcam, 1:800, cat. # AB113); anti-LMX1A (Sigma-Aldrich, 1:400, cat. # AB10533); donkey anti-mouse IgG (Invitrogen, Alexa Fluor 488, 1:500, cat. # A21202); donkey anti-mouse IgG (Invitrogen, Alexa Fluor 647, 1:500, cat. # A31571); donkey anti-mouse IgG (Invitrogen, Alexa Fluor 568, 1:500, cat. # A10037); donkey anti-goat IgG (Invitrogen, Alexa Fluor 647, 1:500, cat. # A21447); donkey anti-sheep IgG (Invitrogen, Alexa Fluor 647, 1:500, cat. # A21448); donkey anti-rabbit IgG (Invitrogen, Alexa Fluor 568, 1:500, cat. # A10042); donkey anti-rabbit IgG (Invitrogen, Alexa Fluor 488, 1:500, cat. # A21206); donkey anti-rabbit IgG (Invitrogen, Alexa Fluor 647, 1:500, cat. # A31573); and goat anti-rat IgG (Invitrogen, Alexa Fluor 488, 1:500, cat. # A48262).

### Microscopy imaging acquisition and quantitative analysis

Brightfield and polarized light images of cortical organoid sections were acquired using an Olympus VS200 slide scanner (Evident Scientific) equipped with a 20× objective. Images were collected for morphological assessment and visualization of microplastic particles. Whole-slide immunofluorescence images were acquired using an Olympus VS200 slide scanner (Evident Scientific) equipped with a 20× objective. Additional fluorescence images presented in the supplementary figures were acquired using an EVOS™ M7000 Imaging System (Thermo Fisher Scientific) with a 20× objective. Exposure settings were maintained across all experimental groups within each staining panel to ensure consistent image acquisition. Image analysis was performed using QuPath (version 0.7.0), CellProfiler (version 4.2.8), MATLAB (R2018b, MathWorks), and GraphPad Prism (version 11; GraphPad Software). Entire organoid sections were manually annotated in QuPath, and annotated regions were exported as individual TIFF images using a custom QuPath script developed in-house for downstream analysis. Exported images were analyzed using CellProfiler, where nuclei were segmented and marker-positive cells were identified using manually determined fluorescence intensity thresholds that were applied uniformly to all images within each staining experiment. Total nuclei count, marker-positive cell counts, and the percentage of marker-positive cells were quantified for each tissue section. CellProfiler output was subsequently processed using a custom MATLAB (R2018b) script developed in-house to summarize quantitative measurements on a per-image basis. Statistical analyses and graphical visualizations were performed using GraphPad Prism (version 11).

### Statistical analysis

Statistical analyses were performed using GraphPad Prism (version 11; GraphPad Software). Statistical tests, sample sizes, and definitions of biological replicates are described in the corresponding figure panels and/or legends for each experiment. Data are presented as indicated in the individual figure panels. Statistical significance was determined using the statistical tests specified for each analysis, with *P* < 0.05 considered statistically significant.

### RNA extraction and bulk RNA sequencing

Total RNA was extracted from cortical organoids using the RNeasy Mini Kit (Qiagen, cat. #74104). Organoids were transferred to 1.5-ml microcentrifuge tubes, washed twice with PBS, and RNA-extracted according to the manufacturer’s instructions. Purified RNA was stored at −80°C. RNA samples were processed by Novogene Co., Ltd. for quality assessment, poly(A)-enriched directional mRNA library preparation, and sequencing on a NovaSeq X Plus platform. All samples passed quality control prior to library preparation. Approximately 9 Gb of raw sequencing data were generated for each sample.

### Bulk RNA sequencing data analysis

Raw sequencing data were used for downstream transcriptomic analyses. Differential gene expression analysis was performed between experimental groups, and genes were ranked according to differential expression statistics. Gene Set Enrichment Analysis (GSEA; Broad Institute) was performed using ranked gene lists to identify significantly enriched biological pathways. Hallmark gene sets from the Molecular Signatures Database (MSigDB) were used as the primary reference gene set collection. Pathways with a false discovery rate (FDR)-adjusted *q*-value < 0.25 were considered significantly enriched, in accordance with GSEA recommendations. Normalized enrichment scores (NES), nominal *P* values, and FDR-adjusted *q* values were used to assess pathway enrichment. Differentially expressed genes and enriched pathways were visualized using volcano plots, heatmaps, enrichment plots, and other graphical representations. From the top 300 gene sets identified in GO analysis as significantly enriched, a total of 67 GO terms representative of phenotypes of interest were selected. A stem plot visualizing −log_10_ FDR q-value with a terminal marker scaled by genes overlapping with the given set (k) was produced using R package ggplot2. For Gene Set Enrichment Analysis (GSEA), the complete set of sequencing-derived genes was ranked in descending order by log_2_ (fold change) values using R to generate a .rnk file. The ranked list was evaluated against the MSigDB gene set collections—specifically Hallmark (h.all.v2026.1.Hs.symbols.gmt), GO Biological Processes (c5.go.bp.v2026.1.Hs.symbols.gmt), and Curated Pathways (c2.all.v2026.1.Hs.symbols.gmt)—using the GSEA desktop application. In parallel, Over-Representation Analysis (ORA) for Hallmark pathways was performed via the GSEA web portal using the subset of significantly upregulated genes. For both approaches, enriched pathways were selected using a significance threshold of *P* < 0.05, and visualized using R.

### Projection of differentially expressed genes onto a whole-body single-cell reference

Two publicly available human single-cell transcriptomic datasets were combined into a single whole-body reference. Whole-body coverage was provided by the Tabula Sapiens “All Cells” object^15^, obtained in H5AD format from the CZ CELLxGENE Discover portal^38^. Cortical coverage at higher cell-type resolution was provided by the Allen Brain Map human middle temporal gyrus (MTG) SMART-seq taxonomy^16^. The Tabula Sapiens object was processed in Python 3.11.15 using Scanpy 1.11.5^39^ and AnnData 0.12.19, read in backed mode so that the full expression matrix was never loaded into memory. Because gene identifiers differed between the two datasets, genes were matched on HGNC symbol. Tabula Sapiens indexes features by Ensembl gene ID. Where several Ensembl identifiers mapped to the same symbol, the first occurrence was retained. To prevent highly sampled organs from dominating the merged reference, cells were subsampled to a maximum of 1,500 per unique tissue × cell-type combination using random sampling with seed 1234. Cells with fewer than 200 detected genes were discarded. Raw counts were exported in Matrix Market format together with the associated cell metadata. Reference assembly and all subsequent analyses were performed in R 4.4.0 (2024-04-24) on x86_64 GNU/Linux (Rocky Linux 8.10; reference BLAS/LAPACK 3.12.0) using Seurat 5.5.1^40^. Allen MTG cells were assigned to major classes by collapsing the subclass_label annotation: excitatory neuron subclasses were assigned to Excitatory neurons; PVALB, SST, VIP, LAMP5 and PAX6 were assigned to Interneurons; Astrocyte was assigned to Astrocytes; Oligodendrocyte was assigned to Oligodendrocytes; OPC was assigned to OPCs; and Endothelial, Pericyte and VLMC were assigned to Endothelial. Tabula Sapiens cells were assigned using the compartment annotation: endothelium to Endothelial, epithelium to Epithelial, immune to Immune, stromal to Stromal, and germline to Germline. Cells belonging to the Tabula Sapiens neural compartment were excluded so that all neural annotation in the merged reference derived from the higher-resolution Allen taxonomy rather than being mixed with a coarser parallel labelling. Cells with an unassigned compartment (none) were also excluded. The two objects were merged, layers joined, and the combined matrix normalized using LogNormalize with a scale factor of 10,000. The 2,000 most variable features were identified by variance-stabilizing transformation, scaled, and used for principal component analysis (50 components). Principal components were computed by truncated SVD using irlba 2.3.7, and a UMAP embedding was computed from the first 30 components using the uwot implementation^41^ (v0.2.4). The final reference comprised 538,999 cells and 50,281 genes. All analyses were performed in parallel at two annotation resolutions. At the major resolution, cells were grouped into six classes: Neural, Immune, Stromal, Epithelial, Endothelial, and Germline, with all Allen-derived cells pooled into a single Neural class. At the fine resolution, 56 classes were used: the 18 Allen subclass_label categories together with the 38 Tabula Sapiens broad_cell_class categories present after compartment filtering. Class labels occurring in both source datasets were disambiguated by prefix to prevent cells of different origin from being pooled. The two resolutions differed only in cell labelling; the underlying reference, gene sets, and principal component embeddings were identical. To determine which cell populations express the genes altered by microplastic exposure, differentially expressed genes (DEGs) were projected onto the single-cell reference following the approach described by Marquez-Galera et al.^42^, originally developed in Cid et al.^43^, with the modifications detailed below. Differential expression between microplastic-exposed and control organoids at day 30 was assessed with DESeq2^44^. Genes were ranked by adjusted *P* value and separated by the sign of the log_2_ fold change. For each direction, the 50 most significant genes also present in the reference were retained. For each DEG set, the reference expression matrix was re-scaled using only those 50 genes and principal component analysis was recomputed on that restricted feature set, yielding a low-dimensional space in which cell position was determined solely by the DEG signature. Reference cells were then binned into four equal-width intervals along PC1 and, separately, along PC2, and the cell-type composition of each bin was computed as the percentage of cells belonging to each class. Pairwise Pearson correlation coefficients were computed between the selected DEGs across all reference cells using log-normalized expression values; genes with zero variance were excluded. Genes were clustered hierarchically on the distance 1 − *r* using average linkage, and co-expression modules were defined with the dynamic tree-cut algorithm^45^(dynamicTreeCut 1.63-1) using deepSplit = 2 and a minimum cluster size of 5. Genes not assigned to a module by the algorithm were reported as unassigned. Dimensionality-reduction plots of the full reference show all 538,999 cells. For t-distributed stochastic neighbor embedding (Rtsne 0.17) and for the DEG-space scatter plots, a class-balanced subsample was used for legibility, with up to 1,200 and up to 800 cells per class, respectively, using seed 1234. This affected visualization only, and all reported compositions, correlations, and modules were computed on the complete reference. Figures were generated with ggplot2 4.0.3, patchwork 1.3.2, cowplot 1.2.0, and pheatmap 1.0.13. Analyses were performed in R 4.4.0 (2024-04-24) on x86_64 GNU/Linux (Rocky Linux 8.10; reference BLAS and LAPACK 3.12.0) with Seurat 5.5.1, SeuratObject 5.4.0, Matrix 1.7-5, irlba 2.3.7, RSpectra 0.16-2, uwot 0.2.4, Rtsne 0.17, dynamicTreeCut 1.63-1, pheatmap 1.0.13, ggplot2 4.0.3, patchwork 1.3.2, cowplot 1.2.0, data.table 1.18.4, dplyr 1.2.1, tidyr 1.3.2, tibble 3.3.1, stringr 1.6.0, and purrr 1.2.2. Pre-processing of the Tabula Sapiens object was performed in Python 3.11.15 with Scanpy 1.11.5, AnnData 0.12.19, h5py 3.16.0 (HDF5 2.1.0), SciPy 1.17.1, NumPy 2.4.6, and pandas 2.3.3. Analyses were run on the Alpine high-performance computing cluster at the University of Colorado Boulder. Reference construction was performed on a single high-memory node with 500 GB of RAM (peak resident memory 198 GB); the projection analyses were run on standard compute nodes. Intermediate and final objects were stored on the CU Boulder PetaLibrary^46^.

### Single-nuclei RNA sequencing

Cortical organoids were transferred to sterile 2-ml microcentrifuge tubes, and culture medium was completely removed. Organoids were washed once with PBS, excess PBS was carefully aspirated, and samples were immediately flash-frozen on dry ice. Frozen samples were stored at −80°C until processing. Samples were subsequently submitted to Novogene Co., Ltd. for tissue dissociation, nuclei isolation, library preparation, and sequencing. Single-nuclei libraries were prepared using the Chromium GEM-X Single Cell 3′ Gene Expression v4 platform (10x Genomics). Libraries were sequenced on an Ultima UG100 sequencing platform. Library quality was assessed using Qubit fluorometric quantification, fragment-size analysis, and quantitative PCR prior to sequencing.

### Single-nuclei RNA sequencing data analysis

Reads were processed with Cell Ranger 10.0.0 (10x Genomics) against the GRCh38-2024-A human reference transcriptome with intronic reads included, as appropriate for nuclear RNA. One library was sequenced per condition (Untreated, sMNP, eMNP). Cell Ranger called 13,285 (Untreated), 15,237 (sMNP) and 17,483 (eMNP) nuclei, with mean read depths of 31,452, 25,590 and 22,044 reads per nucleus, respectively. Only the filtered feature-barcode matrices were used downstream.

For quality control: All subsequent processing was done in R (4.4.3 on the University of Colorado Alpine cluster, 4.5.2 locally) with Seurat 5.5.1 and SeuratObject 5.4.0. Each sample was processed separately. Nuclei were retained when they had at least 1,700 UMIs and 1,000 detected genes (Untreated and eMNP) or 1,300 UMIs and 900 genes (sMNP), a UMI-to-gene ratio of at least 1.2, and at most 25% of UMIs from mitochondrial genes (MT-). No cluster was removed at this stage.

Doublets were identified per sample with scDblFinder (version 1.23.4, seed 42) in cluster-informed mode, using clusters obtained from the SCTransform-normalised data (2,000 variable features, 20 principal components, shared-nearest-neighbour graph with k = 25, Louvain resolution 0.7); the expected doublet rate was estimated by scDblFinder from the data. Nuclei called doublets were removed. Cell-cycle phase was scored with Seurat’s CellCycleScoring (2019 gene lists) for diagnostic purposes only. After quality control and doublet removal, 8,824 (Untreated), 11,067 (sMNP) and 12,215 (eMNP) nuclei remained. For normalisation, dimensionality reduction and choice of clustering parameters, each sample was normalised with SCTransform (sctransform 0.4.3; 2,000 variable features, no covariates regressed), followed by PCA and UMAP (20 principal components; n.neighbors = 10 for the embeddings used for clustering decisions). Clustering used Seurat’s shared-nearest-neighbour graph and the Louvain algorithm.

Clustering parameters were chosen on the Untreated sample and then held fixed for all samples. Every combination of neighbourhood size k (10, 25, 50, 75) and resolution (0.2 to 1.6 in steps of 0.2) was scored on (i) the mean silhouette width in PCA space, (ii) neighbourhood purity, (iii) stability as the mean adjusted Rand index between the clustering of the full data and of 20 bootstrap resamples, and (iv) marker support, defined as the number of clusters lacking at least three genes with AUROC ≥ 0.70 against their most similar cluster. Combinations with stability < 0.80, more than 20% of clusters without marker support, or more than one cluster smaller than 20 nuclei were rejected; among the remainder the finest partition was taken. This selected k = 50 and resolution 1.0 (bootstrap stability 0.833), giving 16 clusters in the Untreated sample.

The 16 Untreated clusters were annotated from three lines of evidence produced by the deposited pipeline (Python 3.11.15; scanpy 1.11.5, scvi-tools 1.4.2, scArches 0.6.1, HNOCA-tools 0.2.1 including snapseed).

#### Reference mapping

Raw counts of the 8,824 Untreated nuclei were mapped with the HNOCA-tools AtlasMapper onto two references. The primary reference was the Human Neural Organoid Cell Atlas (HNOCA; He, Dony, Fleck et al. 2024) restricted to guided cortical protocols (organ annotated as cerebral cortex or telencephalon) at 20–50 days of differentiation, matching the age of the query (day 36), and subsampled to at most 3,000 cells per annot_level_2 label (27,800 cells). Counts were taken from the atlas’s published length-normalised count layer. Query nuclei were embedded into the atlas latent space by training scANVI de novo on the reference (30 latent dimensions, 2 layers, 200 scVI and 20 scANVI epochs) and mapping the query by scArches surgery (100 query epochs). Labels were transferred by weighted k-nearest neighbours (k = 100) using 3,000 highly variable genes shared between query and reference. The developing human brain atlas of Braun et al. (2023; Class level, subsampled to 1,000 cells per label, 100,000 cells maximum; mapped with the same scANVI procedure) served as a complementary reference to flag possible off-target populations; it was not used to assign identities. Presence scores were computed to identify reference populations absent from the organoids.

#### Marker-based scoring

Independently, clusters were scored with snapseed (HNOCA-tools) against a hierarchical marker list ordered lineage and differentiation state → cortical identity → regional identity, using positive and negative markers, with descent to a finer label only when the best child scored ≥ 0.10 above chance and ≥ 0.15 above the runner-up.

#### Differential expression between clusters

One-versus-rest differential expression was computed per cluster on log-normalised counts with the Wilcoxon rank-sum test (scanpy rank_genes_groups, tie-corrected, Benjamini–Hochberg adjustment); candidate markers were ranked by the difference in the fraction of expressing nuclei, requiring adjusted P < 0.05, log_2_ fold change ≥ 0.5 and detection in ≥ 20% of the cluster’s nuclei (Supplementary Table 1). Because clusters were defined from the same data, these P values were used only to rank genes. Expression of canonical markers of the developing human cortex was additionally inspected per cluster as log_2_(counts per 10,000 + 1).

Final labels were assigned by the authors from the combined evidence; 16 clusters: dorsal radial glia (two clusters), dorsal progenitors, roof plate-like/dorsal progenitors, S-phase progenitors, G2/M progenitors, stress-responsive cortical radial glia, cortical neural progenitors, neurogenic intermediate progenitor cells, maturing glutamatergic neurons (two clusters), maturing cortical neurons, dorsal sensory/interneuron-like neurons, LHX9/MEIS2 neurons, PAX2/LHX1 GABAergic neurons and mesenchymal cells). At day 36 many clusters represent successive maturation states rather than distinct cell types, and labels were kept at that level of description.

Subsequently, the quality-controlled nuclei of the three conditions (32,106 in total) were merged without integration, log-normalised (scale factor 10,000), and embedded together (3,000 variable features, scaled data, PCA, UMAP on 20 principal components with n.neighbors = 10, min.dist = 0.3, seed 42).

The merged data were clustered over a grid of k (20, 30, 50, 80) and resolution (0.2, 0.4, 0.6, 0.8, 1.0, 1.2, 1.6) on the 20 principal components. Each option was scored on the Untreated nuclei only against the 16-cluster Untreated partition, together with per-cluster sample mixing entropy and cluster sizes, and the options were inspected visually against the Untreated clusters and annotation drawn on the same embedding. The selected clustering (k = 80, resolution 0.8) gave 20 clusters and had the highest agreement with the Untreated partition. Merged clusters are reported by number. Their identity is given by the Untreated nuclei they contain.

Composition was analyzed separately for each comparison (Untreated versus sMNP, Untreated versus eMNP) on the nuclei of the two conditions concerned. For each merged cluster the proportion of nuclei in each condition is reported with a 95% Wilson score interval, which describes the sampling precision of the proportion within that library. Because each condition was captured once, these intervals do not capture between-organoid-batch variability, and no hypothesis test of the treatment effect on composition was performed.

Differential expression between the treated and Untreated nuclei was computed with Seurat’s FindMarkers (two-sided Wilcoxon rank-sum test on log-normalised counts), for all nuclei together and within each merged cluster with at least 30 nuclei per condition. Genes detected in fewer than 10 nuclei, or in fewer than 10% of nuclei in both groups, were not tested. P values were FDR-corrected over the genes tested in each contrast; genes with adjusted P < 0.05 and |log_2_ fold change| ≥ 0.25 were called differentially expressed (Supplementary Tables 2 and 3).

Cell Ranger 10.0.0; R 4.4.3 (Alpine) and 4.5.2; Seurat 5.5.1; SeuratObject 5.4.0; sctransform 0.4.3; scDblFinder 1.23.4; presto 1.1.0; ggplot2 4.0.3; Python 3.11.15 with scanpy 1.11.5, anndata 0.10.8 and 0.12.19, scvi-tools 1.1.6 and 1.4.2, scArches 0.6.1, HNOCA-tools 0.2.1 (snapseed), PyTorch 2.5.1. All code, configuration files and the cluster label tables are available at https://github.com/aguado-lab/2026-microplastics (single-cell/), together with the SLURM resource specifications used on Alpine and instructions to run each stage on a workstation.

## Data availability

Data have been submitted to GEO and are available upon reasonable request.

## Code availability

Code used to analyze sequencing data is available on GitHub, which is available upon request. Custom QuPath and MATLAB scripts used for image export and quantitative analysis and R scripts used for Bulk RNA sequencing analysis are available from the corresponding author upon reasonable request.

## Contributions

C.P., and D.M. generated human brain organoids. R.K. and M.G. prepared the sMNPs and eMNPs, respectively. C.P., M.S-C., R.K., M.G., D.M., R.M., C.M., M.C., A.H., A.C-M., and J.A. contributed to acquisition, analysis or interpretation of data. C.P., D.M., R.M. and M.S-C. analyzed transcriptomic data. J.A., C.P., R.K., and A.H. contributed to experimental design. C.P. and J.A. wrote the paper. J.A. planned and supervised the project. All authors edited and approved the final version of this article.

## Acknowledgements

We thank the University of Colorado Anschutz Advanced Light Microscopy Core for technical support; Drs Monica Serban and Elizabeth Arrigali at the University of Montana for the use of equipment and instruction; the University of Montana Raman, Confocal, and Hyperspectral-darkfield Microscopy Core for access to instrumentation and technical assistance; and J.A. laboratory members for discussions. This work utilized the Alpine High-Performance Computing resource at the University of Colorado Boulder. Alpine is jointly funded by the University of Colorado Boulder, the University of Colorado Anschutz, Colorado State University, and the National Science Foundation (award 2201538). Data storage is supported by the University of Colorado Boulder "PetaLibrary". M.A.G. was supported by K08 (ES038232) and K12 (TR005467) NIH awards. M.J.C. was supported by a U01 (AG08855) NIH award. A.C-M. was supported by an R01 (7R01MH129732-04) award and startup funding from the Department of Anesthesiology and the NeuroTechnology Center at the University of Colorado Anschutz Medical Campus. A.H. and J.A. were supported by a joint Skaggs Scholars program grant for the study of microplastics’ effects in the nervous system. J.A. was supported by startup funding from the Department of Pharmaceutical Sciences at the University of Colorado Anschutz Medical Campus. The funders had no role in study design, data collection and analysis, decision to publish or preparation of the manuscript.

## Supplementary Figure Legends

**Supplementary figure 1:**
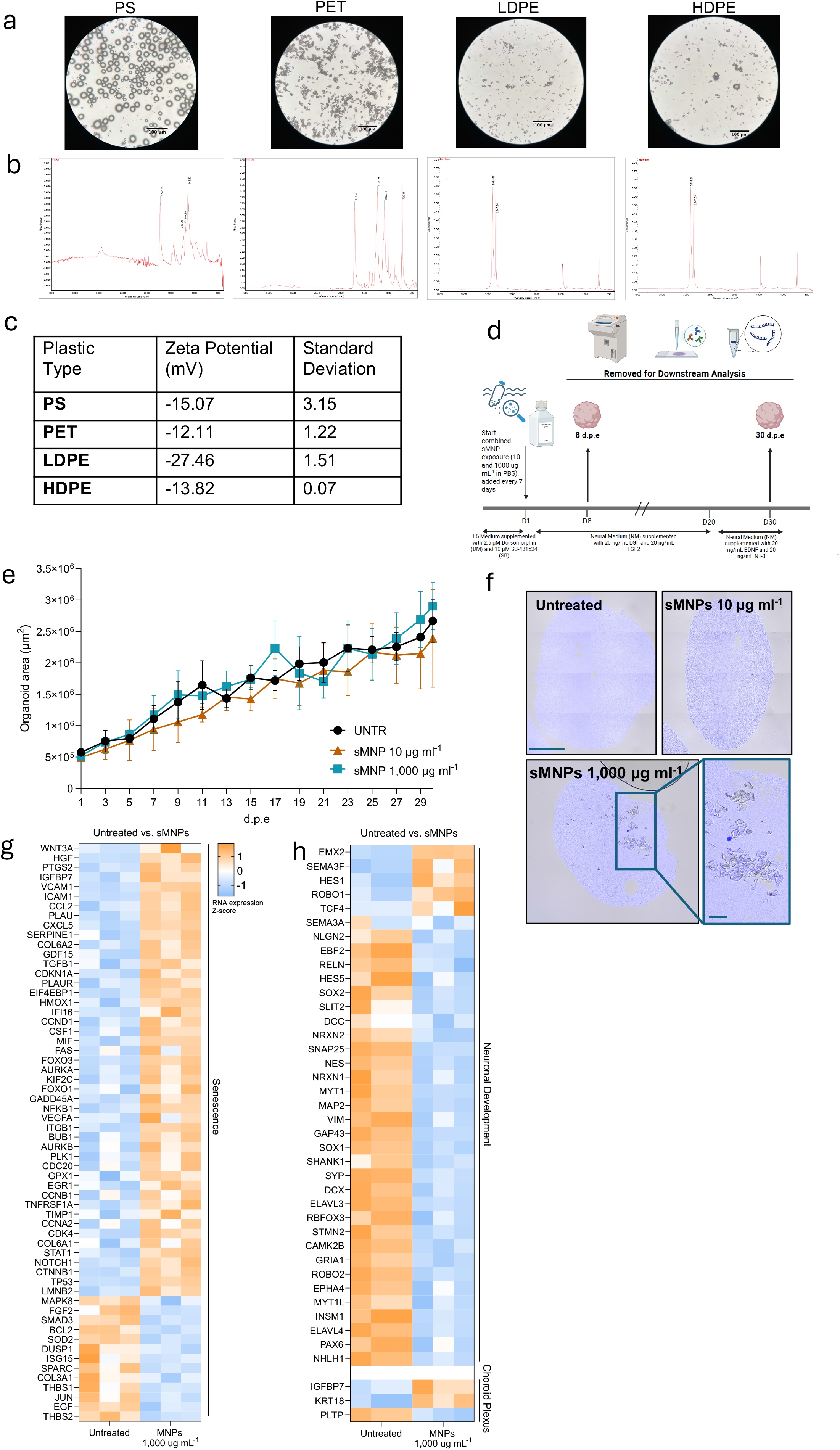
Synthetic MNP characterization, experimental design, and longitudinal cortical organoid growth. (**a**) Brightfield images taken in the 40X objective for each sMNP used for CO exposure: polystyrene (PS), polyethylene terephthalate (PET), low-density polyethylene (LDPE), and high-density polyethylene (HDPE). (**b**) FTIR confirmation of each sMNP composition. Plastic composition was confirmed using ATR-FTIR. Spectra were compared to known peaks for respective polymers to confirm microplastic structure. From left to right: FTIR spectra for PS, PET, LDPE, HDPE. (**c**) Zeta potential (mV) of PS, PET, LDPE, and HDPE microplastic suspensions measured by electrophoretic light scattering using a Malvern ZetaSizer Ultra. Microplastics were dispersed in MilliQ water containing 0.05% Tween 20 and sonicated prior to analysis. Values are presented as mean ± SD. (**d**) Schematic overview of experimental design. Human iPSC-derived COs were exposed to sMNPs (10 or 1,000 μg ml^-1^ in PBS) beginning on day 1, with sMNPs replenished every 7 days. COs were cultured in differentiation media supplemented as indicated throughout the exposure period. Samples were collected after 8 d.p.e. or 30 d.p.e. for downstream analysis. Created with BioRender. (**e**) Size quantification of untreated, 10 μg ml^-1^ and 1,000 μg ml^-1^ sMNP-treated COs over 30 days of exposure. Data are presented as mean ± SD. *n* = 5-19 organoids per treatment group from a single organoid differentiation. (**f**) Representative fluorescence images of DAPI-stained untreated, 10 μg ml^-1^ and 1,000 μg ml^-1^ sMNP-treated COs, exposed on day 6 of growth and collected 30 d.p.e. The boxed region in the 1,000 μg ml^-1^ condition is shown at higher magnification to highlight particle aggregates. Scale bars = 200 μm (lower magnification) and 50 μm (higher magnification). (**g**) Heatmap of selected senescence associated genes identified from the SenSig^13^ and SenMayo^14^ gene sets in untreated and 1,000 μg ml^-1^ sMNP-treated COs, exposed on day 6 of growth and collected 8 d.p.e. Gene expression is displayed as row-wise Z-scores of normalized RNA-sequencing data, with orange representing higher relative expression and blue representing lower relative expression. (**h**) Heatmap of selected neuronal development and choroid plexus marker genes in untreated and 1,000 μg ml^-1^ sMNP treated COs, exposed on day 6 of growth and collected 8 d.p.e. Expression values are shown as row-wise Z-scores of normalized RNA-sequencing data, with orange representing higher relative expression and blue representing lower relative expression. Genes are grouped according to their annotated biological function.

**Supplementary figure 2:**
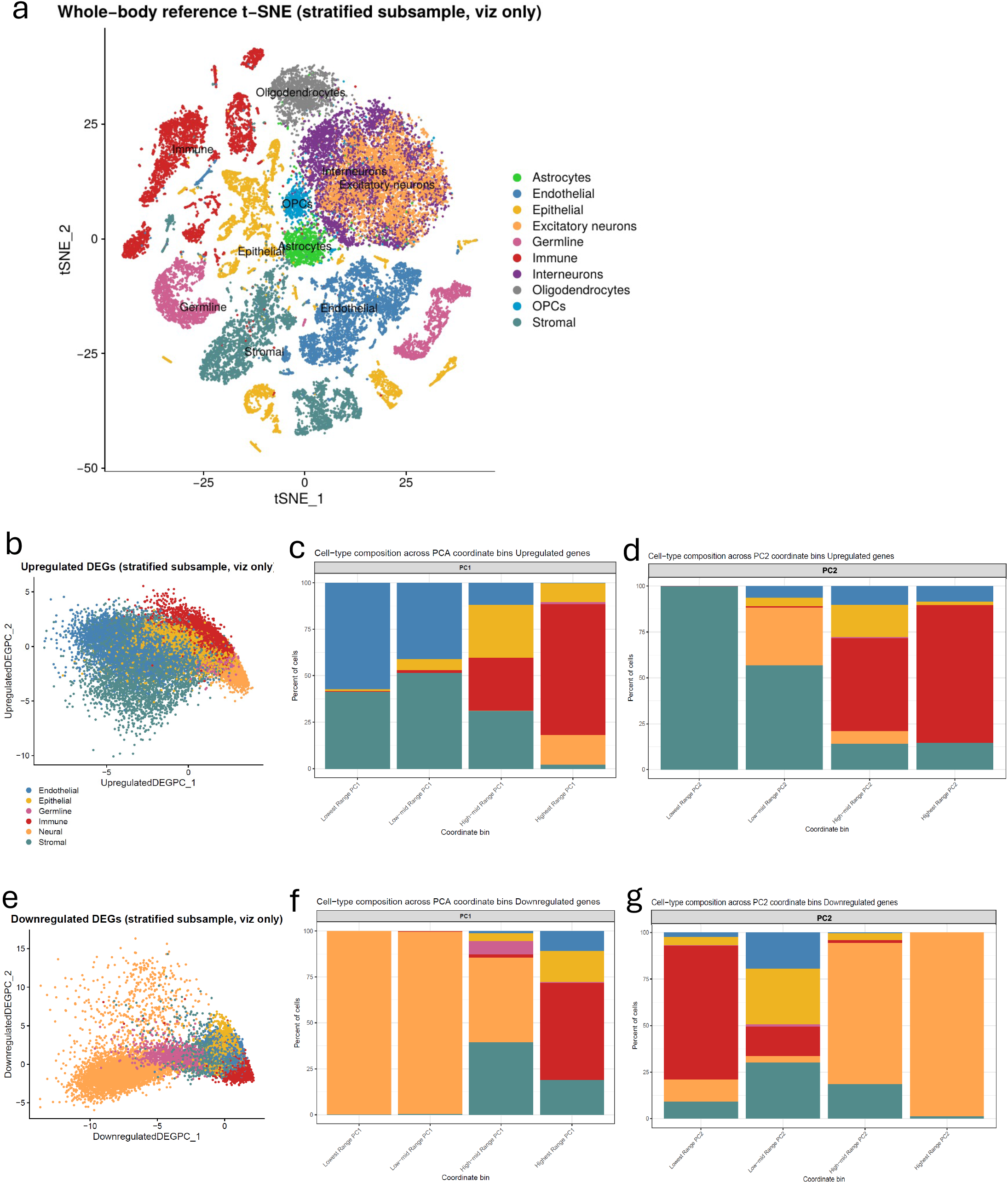
Low-resolution cell-type projection of sMNP-associated transcriptional signatures in cortical organoids. (**a**) t-SNE visualization of the whole body reference showing ten cell classes, including five brain-derived populations (excitatory neurons, interneurons, astrocytes, oligodendrocytes, and OPCs) and five broader whole-body classes (immune, epithelial, endothelial, stromal, and germline). Brain-derived populations were subsequently pooled into a single neural class for DEG projection analyses. (**b**) PCA visualization of single-cell reference populations based on the top 50 upregulated DEGs in cortical organoids exposed to sMNPs on day 6 of growth and collected 30 d.p.e., ranked by adjusted *P* value. Reference cells were derived from a merged Tabula Sapiens and Allen Brain Map dataset and colored according to cell class^15,16^. PCA plots display a class-balanced, stratified subsample of reference cells for visualization. **c,d,** Cell-type composition across PCA coordinate bins for PC1 (**c**) and PC2 (**e**) of the upregulated DEG signature in COs exposed to sMNPs on day 6 of growth and collected 30 d.p.e. Stacked bars represent the percentage of reference cells assigned to endothelial, epithelial, germline, immune, neural, and stromal cell classes within each PCA coordinate bin. (**e**) PCA visualization of single cell reference populations based on the top 50 downregulated DEGs in cortical organoids exposed to sMNPs on day 6 of growth and collected 30 d.p.e., ranked by adjusted *P* value. Reference cells were derived from a merged Tabula Sapiens and Allen Brain Map dataset and colored according to cell class^15,16^. PCA plots display a class-balanced, stratified subsample of reference cells for visualization. **f,g,** Cell-type composition across PCA coordinate bins for PC1 (**f**) and PC2 (**g**) of the downregulated DEG signature in COs exposed to sMNPs on day 6 of growth and collected 30 d.p.e. Stacked bars represent the percentage of reference cells assigned to endothelial, epithelial, germline, immune, neural, and stromal cell classes within each PCA coordinate bin.

**Supplementary figure 3:**
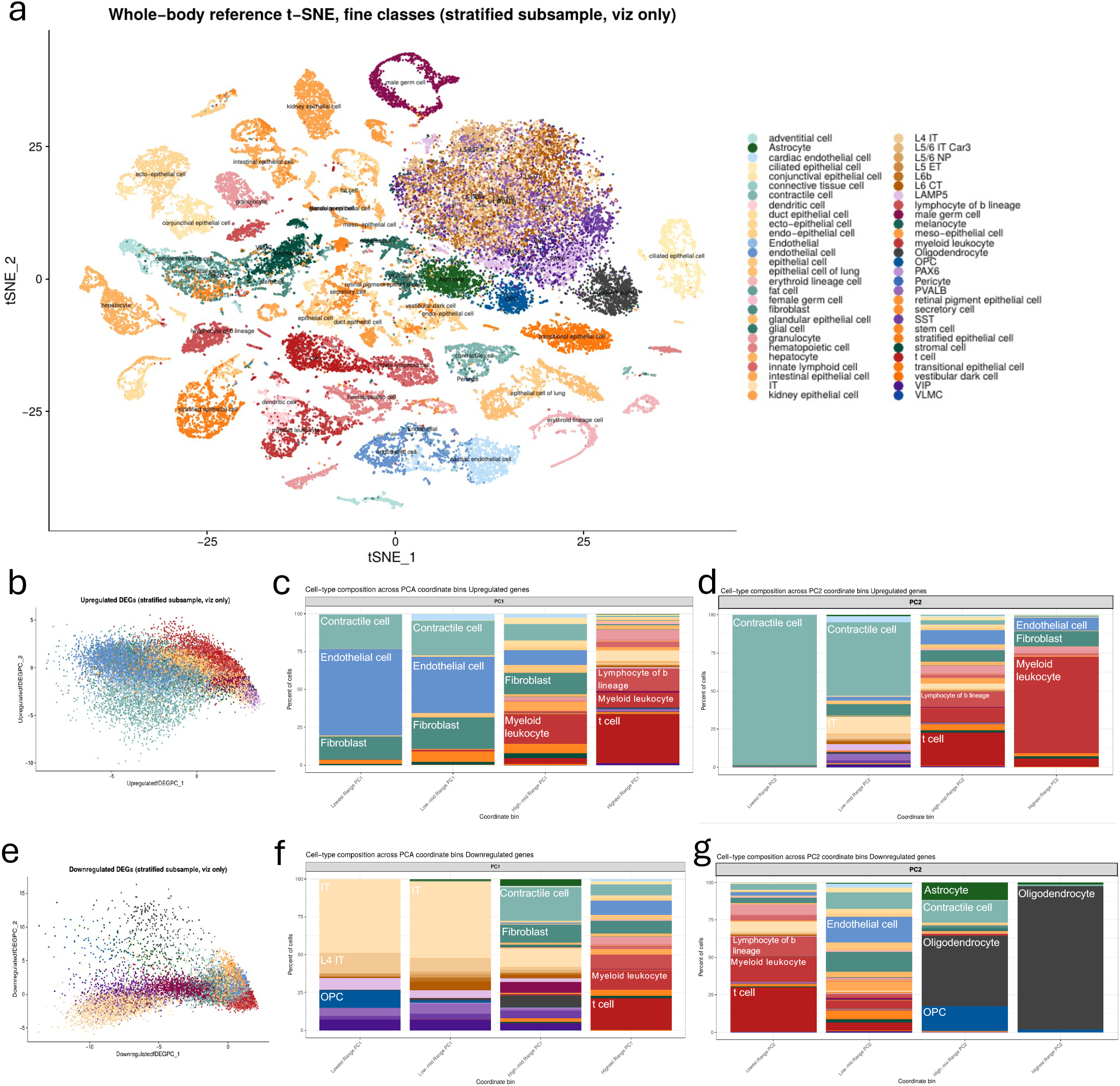
High-resolution cell-type projection of sMNP-associated transcriptional signatures in cortical organoids. (**a**) t-SNE visualization of the whole-body reference at fine resolution, comprising 56 cell-type categories from the Allen Brain Map and Tabula Sapiens references^15,16^. Brain-derived populations remain individually annotated at this stage. The color codes apply for the entire figure. (**b**) PCA visualization of reference cells based on the top 50 upregulated DEGs in cortical organoids exposed to sMNPs on day 6 of growth and collected 30 d.p.e., ranked by adjusted *P* value. **c,d,** Cell-type composition across PCA coordinate bins for PC1 (**c**) and PC2 (**d**) of the upregulated DEG signature in COs exposed to sMNPs on day 6 of growth and collected 30 d.p.e. Stacked bars represent the percentage of reference cells assigned to each fine-resolution cell type within each PCA coordinate bin. Selected cell-type groups are labeled within the composition plots to facilitate visualization of the predominant populations. PCA projections and cell-type compositions were generated using a stratified subsample for visualization. (**e**) PCA visualization of reference cells based on the top 50 downregulated DEG signatures in cortical organoids exposed to sMNPs on day 6 of growth and collected 30 d.p.e., ranked by adjusted *P* value. **f,g,** Cell-type composition across PCA coordinate bins for PC1 (**f**) and PC2 (**g**) of the downregulated DEG signature. Stacked bars represent the percentage of reference cells assigned to each fine-resolution cell type within each PCA coordinate bin. Selected cell-type groups are labeled within the composition plots to facilitate visualization of the predominant populations. PCA projections and cell-type compositions were generated using a stratified subsample for visualization.

**Supplementary figure 4:**
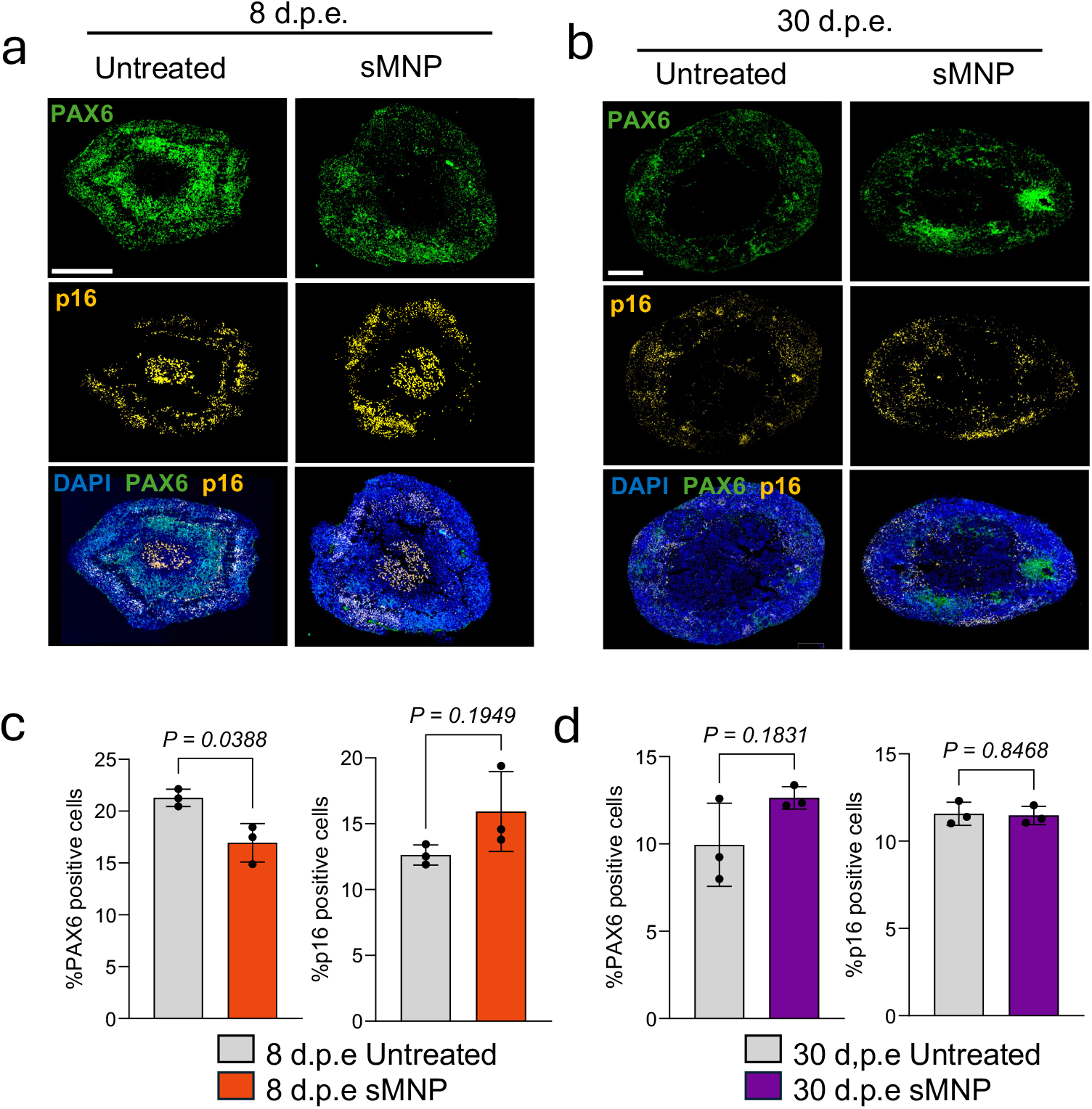
sMNP exposure alters senescence-associated and neurodevelopmental transcriptional signatures in cortical organoids. **a,b,** Representative immunofluorescence images of untreated and 1,000 μg ml^-1^ sMNP treated COs, exposed on day 6 of growth and collected 8 d.p.e. (**a**) and 30 d.p.e. (**b**). Sections were stained for PAX6 (green), and p16 (yellow). Nuclei were counterstained with DAPI (blue). Merged images are shown for each staining panel. Scale bars = 200 μm for 8 d.p.e. images and 250 μm for 30 d.p.e. images. **c,d,** Quantification of PAX6 positive and p16 positive cells in untreated and 1,000 μg ml^-1^ sMNP treated COs, exposed on day 6 of growth and collected 8 d.p.e. (**c**) and 30 d.p.e. (**d**). Data are presented as mean ± SD, with each dot representing an individual organoid. Statistical significance was determined using a two-tailed Welch’s *t*-test. *n*=3 organoids per group. Exact *P* values are shown.

**Supplementary figure 5:**
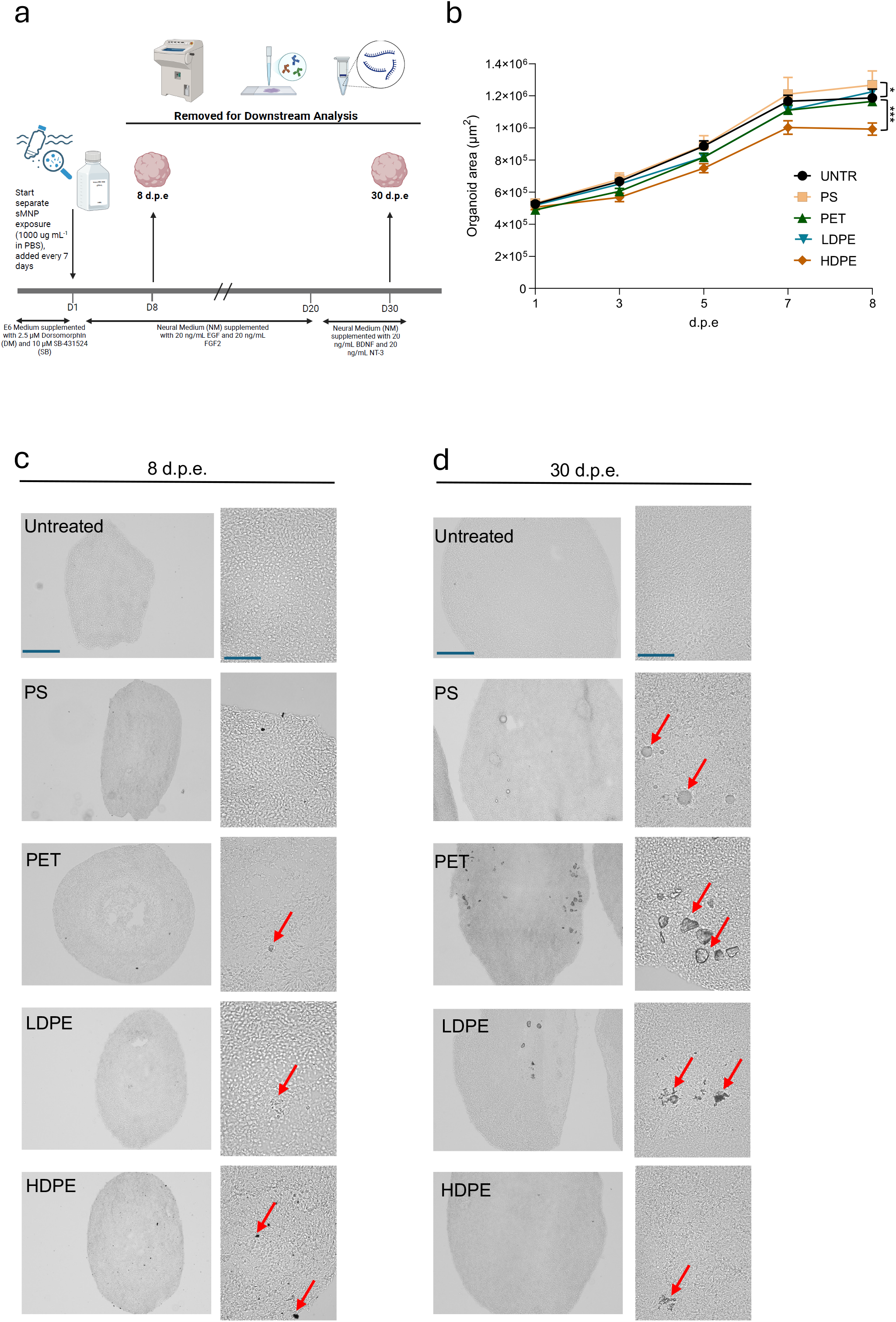
Experimental design and characterization of cortical organoid growth following polymer-specific microplastic exposure. (**a**) Schematic overview of experimental design. Human iPSC-derived COs were treated with individual polymers at 1,000 μg ml^-1^ in PBS, beginning on day 1, with polymers replenished every 7 days. COs were cultured in differentiation media supplemented as indicated throughout the exposure period. Samples were collected after 8 d.p.e. and 30 d.p.e. for downstream analysis. Created with BioRender. (**b**) Size quantification of untreated and 1,000 μg ml^-1^ PS, PET, LDPE, or HDPE-treated COs over 8 days of exposure. Data are presented as mean ± SD. Significance was determined by *p*-value with ***=*p*<0.0001, *=*p*<0.05. *n* = 6-14 organoids per treatment group from a single organoid differentiation. **c,d,** Brightfield images of COs exposed to PS, PET, LDPE or HDPE on day 6 of growth and collected 8 d.p.e. (**c**) and 30 d.p.e. (**d**). Low-magnification images (left) show overall organoid morphology, while higher-magnification images (right) highlight the organoid surface and the presence of associated sMNP particles, indicated by red arrows. Images are representative of independent organoids from each treatment group. Scale bars = 200 μm for lower magnification and 50 μm for higher magnification.

**Supplementary figure 6:**
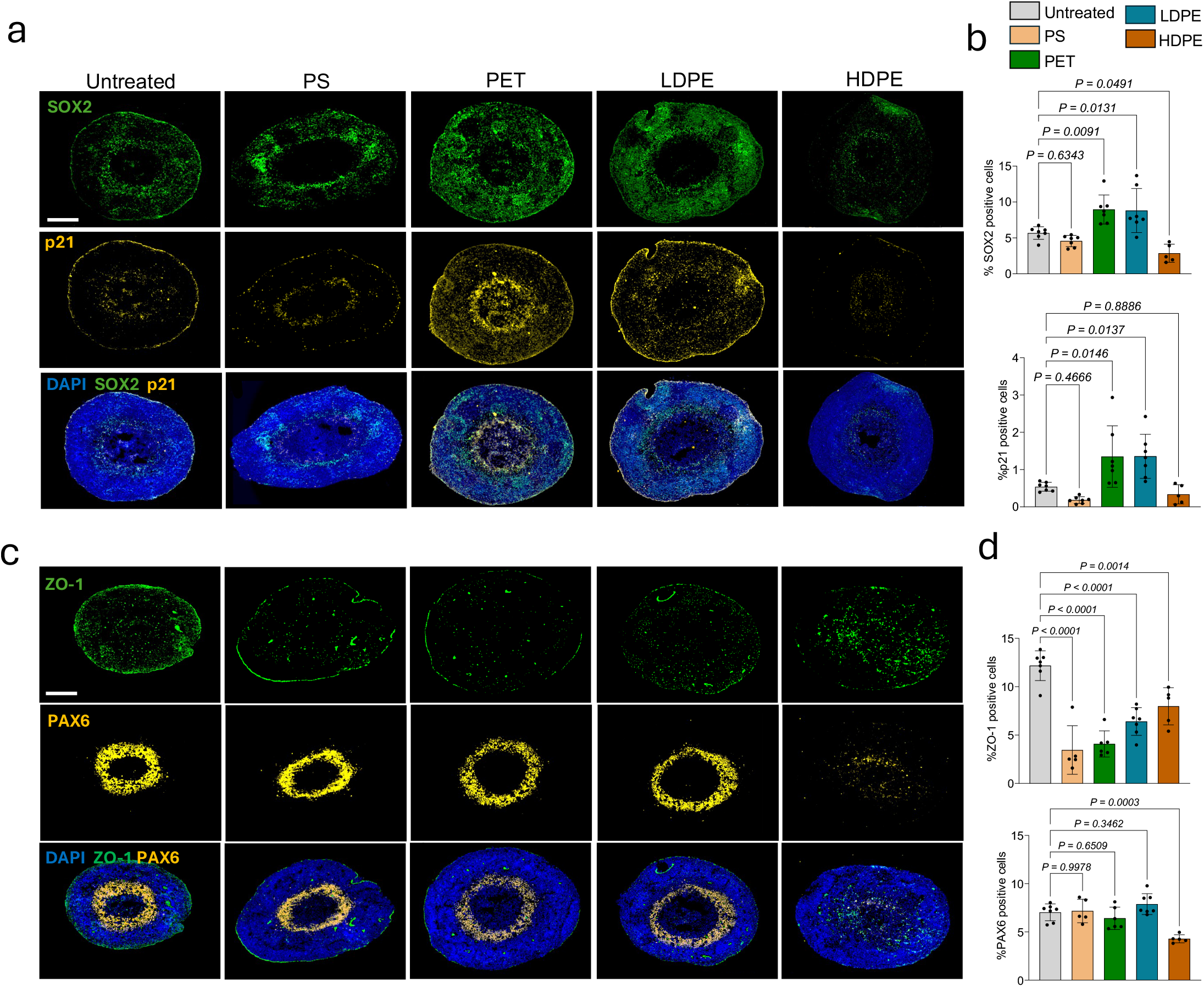
Polymer-specific effects on neural progenitor, neuronal, and senescence-associated markers in cortical organoids. **a,c,e,** Representative immunofluorescence images of untreated and 1,000 μg ml^-1^ PS, PET, LDPE, or HDPE-treated COs, exposed on day 6 of growth and collected 30 d.p.e. Sections were stained for SOX2 (green) and p21 (yellow) (**a**); MAP2 (yellow) (**c**); PAX6 (green) and p16 (yellow) (**e**). Nuclei were counterstained with DAPI (blue). Merged images are shown for each treatment group. Scale bars = 400 μm for untreated, PS, and LDPE SOX2/p21, all MAP2 images and all PAX6/p16 images; 250 μm for PET and HDPE SOX2/p21 images. **b,d,f,** Quantification of SOX2-positive and p21-positive (**b**); MAP2-positive (**d**); PAX6 positive and p16 positive (**f**) cells in untreated and 1,000 μg ml^-1^ PS, PET, LDPE, or HDPE-treated COs, exposed on day 6 of growth and collected 30 d.p.e. Data are presented as mean ± SD, with each data point representing an individual organoid. Data shown are batch-corrected to account for variability between differentiations. Statistical significance was determined using a one-way ANOVA followed by Dunnett’s multiple comparisons test. Exact *P* values are shown. n=4-14 organoids per treatment group.

**Supplementary figure 7:**
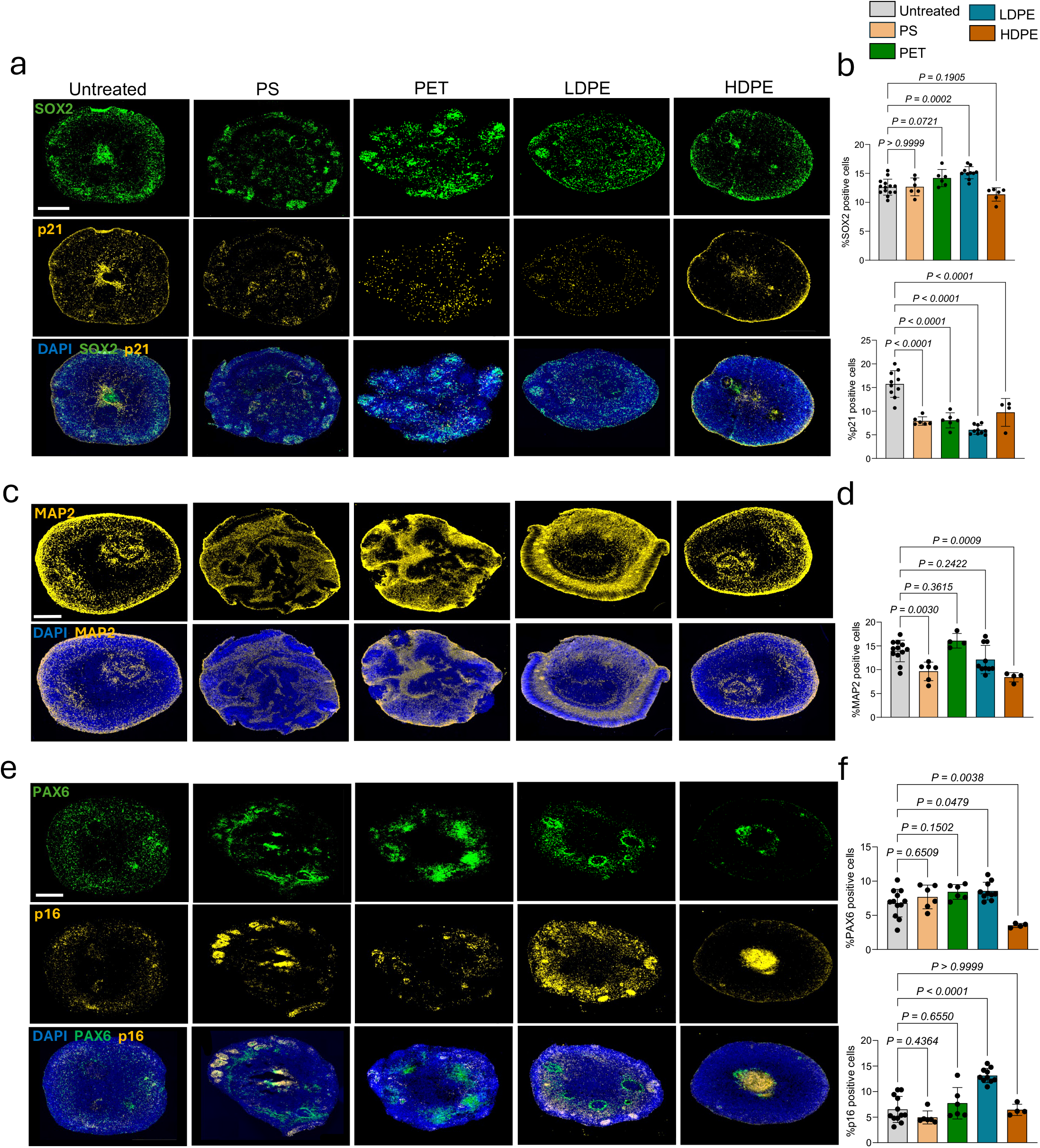
Individual sMNP polymers differentially alter neural progenitor identity, cellular senescence, and tight-junction organization in developing COs. **a,c,** Representative immunofluorescence images of untreated and 1,000 μg ml^-1^ PS, PET, LDPE, or HDPE-treated COs, exposed on day 6 of growth and collected 8 d.p.e. Sections were stained for SOX2 (green) and p21 (yellow) (**a**); ZO-1 (green) and PAX6 (yellow) (**c**). Nuclei were counterstained with DAPI (blue). Merged images are shown for each treatment group. Scale bars = 200 μm. **b,d,** Quantification of SOX2 positive and p21 positive (**b**); ZO-1 positive and PAX6 positive (**d**) cells in untreated and 1,000 μg ml^-1^ PS, PET, LDPE, or HDPE-treated COs exposed on day 6 of growth and collected 8 d.p.e. Data are presented as mean ± SD, with each data point representing an individual organoid. Statistical significance was determined using a one-way ANOVA followed by Dunnett’s multiple comparisons test. Exact *P* values are shown. n = 5-7 organoids per treatment group from a single organoid differentiation.

**Supplementary figure 8:**
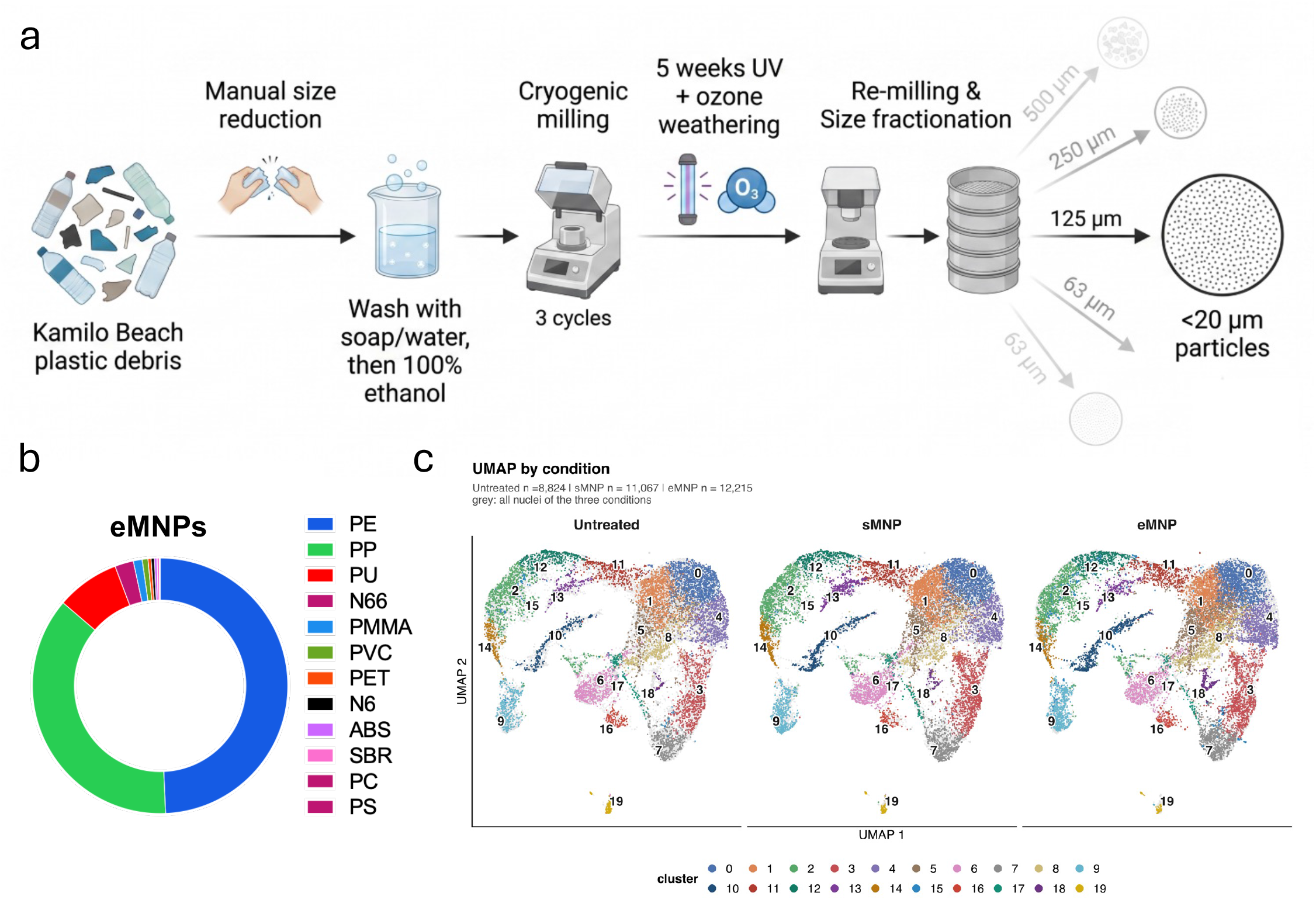
Generation, polymer composition, and transcriptional profiling of environmentally derived micro- and nanoplastics (eMNPs). (**a**) Schematic overview of eMNP generation from plastic debris collected at Kamilo Beach, Hawaii. Plastic debris was manually size-reduced, washed with soap and water followed by 100% ethanol, cryogenically milled for three cycles, and subjected to 5 weeks of UV and ozone weathering. The weathered material was subsequently re-milled and size-fractionated to generate particles of defined size ranges, including 500, 250, 125, 63, and <20 µm, before exposing organoids on day 6 of growth. (**b**) Polymer composition of the resulting eMNP preparation, consisting primarily of polyethylene (PE) and polypropylene (PP), with additional contributions from polyurethane (PU), nylon 66 (N66), polymethyl methacrylate (PMMA), polyvinyl chloride (PVC), polyethylene terephthalate (PET), nylon 6 (N6), acrylonitrile butadiene styrene (ABS), styrene-butadiene rubber (SBR), polycarbonate (PC), and polystyrene (PS). (**c**) UMAP visualization of merged transcriptional clusters across untreated, and 1,000 μg ml^-1^ sMNP or eMNP -treated COs exposed on day 6 of growth and collected 30 d.p.e. Each panel displays nuclei colored by cluster identity, with the same UMAP embedding and cluster assignments shown across conditions. Cluster numbers correspond to the merged cell-state annotations described in the text. Untreated, *n* = 8,824 nuclei; sMNP, *n* = 11,067 nuclei; eMNP, *n* = 12,215 nuclei. Gray points represent nuclei from all three conditions and provide the reference distribution for each condition-specific UMAP.

**Supplementary figure 9:**
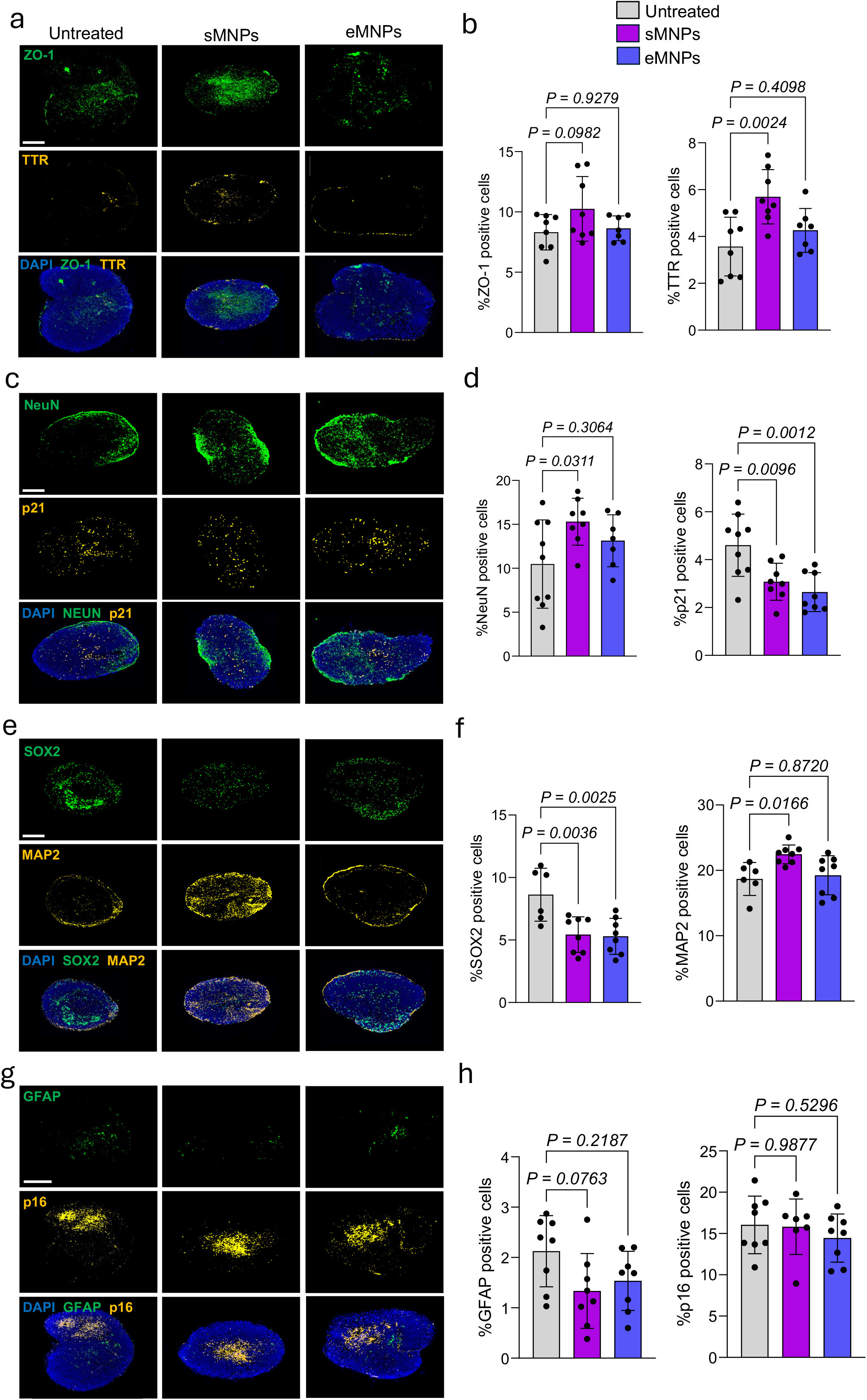
Cellular characterization of cortical organoids following sMNP and eMNP exposure. **a,c,e,g,** Representative immunofluorescence images of untreated, 1,000 μg ml^-1^ sMNP, or eMNP -treated COs exposed on day 6 of growth and collected 30 d.p.e. Sections were stained for ZO-1 (green), and TTR (yellow) (**a**); NeuN (green), and p21 (yellow) (**c**); SOX2 (green), and MAP2 (yellow) (**e**); GFAP (green), and p16 (yellow) (**g**). Nuclei were counterstained with DAPI (blue). Merged images are shown for each staining panel. Scale bars = 200 μm for all images. **b,d,f,h,** Quantification of ZO-1-positive and TTR-positive (**b**); NeuN-positive and p21-positive (**d**); SOX2-positive and MAP2-positive (**f**); GFAP-positive, and p16-positive (**h**) cells in untreated, 1,000 μg ml^-1^ sMNP, or eMNP-treated COs exposed on day 6 of growth and collected 30 d.p.e. Data are presented as mean ± SD, with each data point representing an individual organoid. Statistical significance was determined using a one-way ANOVA followed by Dunnett’s multiple comparisons test. Exact *P* values are shown. n= 5-9 organoids per treatment group from a single organoid differentiation.

**Supplementary figure 10:**
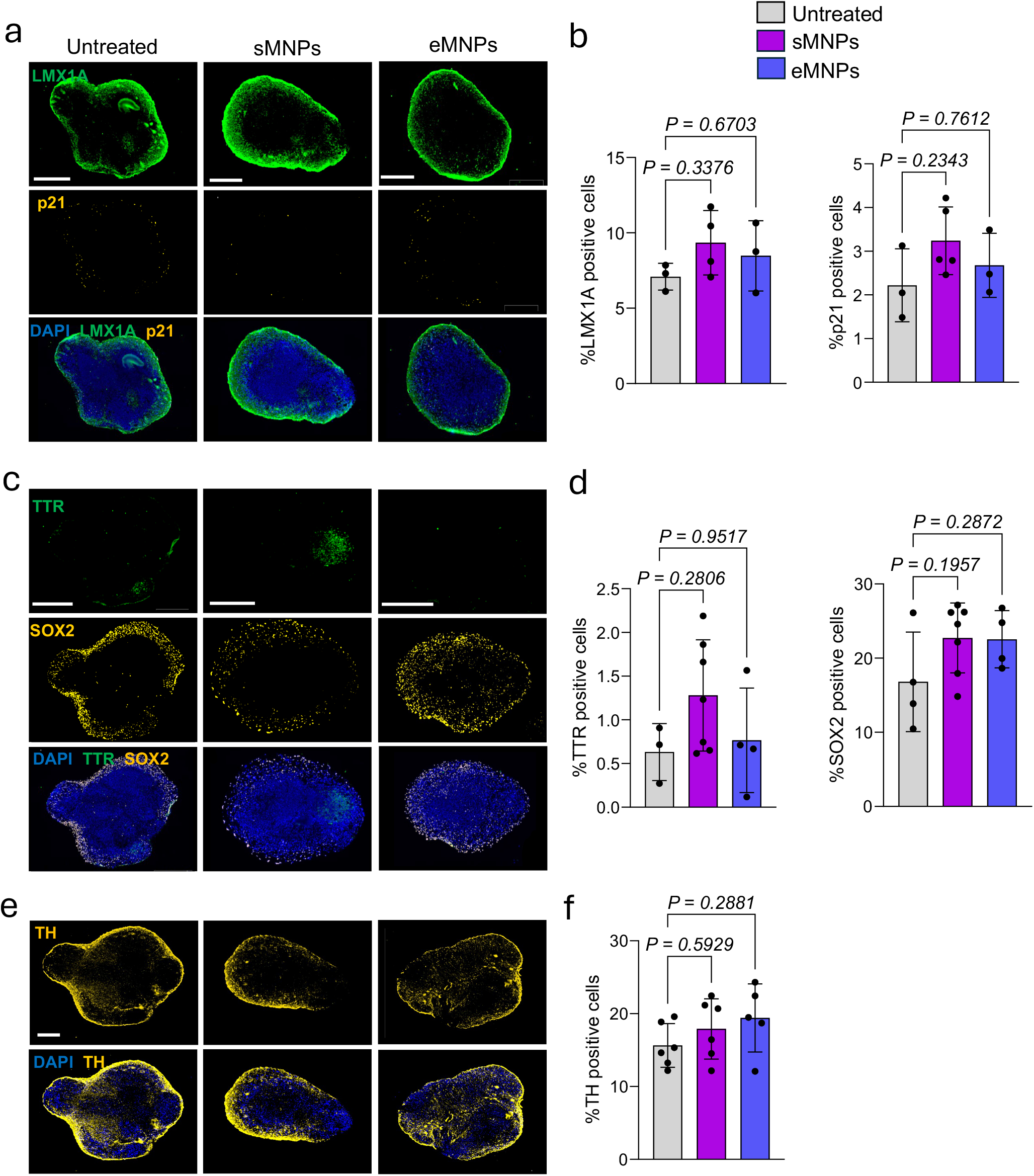
MNP exposure alters cellular phenotypes in mature ventral midbrain organoids. **a,c,e,** Representative immunofluorescence images of untreated, 1,000 μg ml^-1^ sMNP, or eMNP-treated VMOs, exposed on day 90 of growth and collected 30 d.p.e. Sections were stained for LMX1A (green), and p21 (yellow) (**a**); TTR (green), and SOX2 (yellow) (**c**); and TH (yellow) (**e**). Nuclei were counterstained with DAPI (blue). Merged images are shown for each staining panel. Scale bars = 200 μm for GFAP/TH images; 400 μm for untreated and 250 μm for sMNP- and eMNP-treated LMX1A/p21 images; and 400 μm for untreated and eMNP-treated TTR/SOX2 images and 250 μm for sMNPs-treated TTR/SOX2 images. **b,d,f,** Quantification of LMX1A-positive and p21-positive (**b**); TTR-positive, and SOX2-positive (**d**); and TH-positive (**f**) cells in untreated, 1,000 μg ml^-1^ sMNP, or eMNP-treated VMOs, exposed on day 90 of growth and collected 30 d.p.e. Data are presented as mean ± SD, with each data point representing an individual organoid. Statistical significance was determined using a one-way ANOVA followed by Dunnett’s multiple comparisons test. Exact *P* values are shown. n=3-7 organoids per treatment group from a single organoid differentiation.

**Supplementary figure 11:**
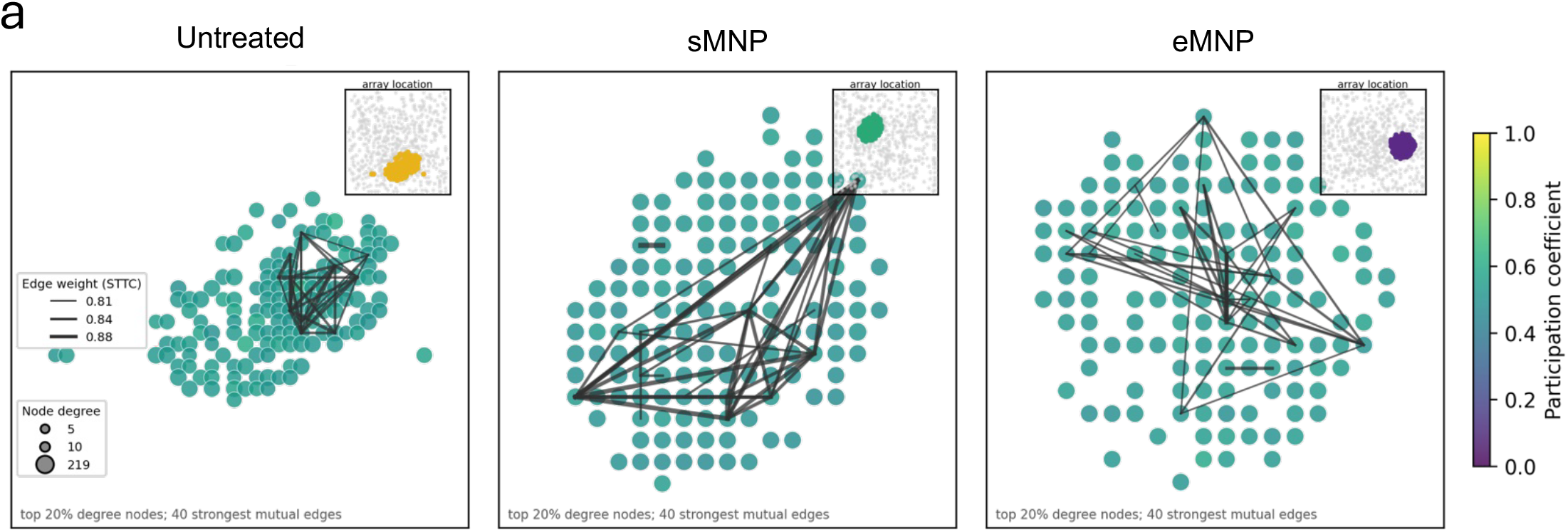
Functional activity and network organization of MNP-exposed ventral midbrain organoids. (**a**) Representative functional networks from an untreated organoid, an sMNP-exposed organoid, and an eMNP-exposed organoid. Each circle represents one of the displayed electrodes, and the connecting lines indicate functional relationships among electrodes based on coordinated spike timing, as measured using the spike-time tiling coefficient (STTC). The panel displays the top 20% of connected electrodes by node degree and up to 40 of their strongest mutual connections. Larger circles indicate electrodes with more functional connections, while thicker lines indicate stronger STTC connectivity. Node color represents the participation coefficient. Values closer to zero indicate predominantly within-module connectivity, whereas higher values indicate connections distributed across multiple modules. The small insets show where the displayed electrodes are located on the complete electrode array.

## References

1 Thompson, R. C. et al. Twenty years of microplastic pollution research-what have we learned? Science 386, eadl2746 (2024). 10.1126/science.adl2746

2 Nihart, A. J. et al. Bioaccumulation of microplastics in decedent human brains. Nat. Med. 31, 1114–1119 (2025). 10.1038/s41591-024-03453-1

3 Runting Li. Microplastics and nanoplastics in brain tumours and the healthy human brain. Nature Health 1, 633–646 (2026). 10.1038/s44360-026-00091-4

4 Chaib, S., Tchkonia, T. & Kirkland, J. L. Cellular senescence and senolytics: the path to the clinic. Nat. Med. 28, 1556–1568 (2022). 10.1038/s41591-022-01923-y

5 Ogrodnik, M. et al. Whole-body senescent cell clearance alleviates age-related brain inflammation and cognitive impairment in mice. Aging Cell 20, e13296 (2021). 10.1111/acel.13296

6 Aguado, J. et al. Senolytic therapy alleviates physiological human brain aging and COVID-19 neuropathology. Nat Aging 3, 1561–1575 (2023). 10.1038/s43587-023-00519-6

7 Zhang, W., Sun, H. S., Wang, X., Dumont, A. S. & Liu, Q. Cellular senescence, DNA damage, and neuroinflammation in the aging brain. Trends Neurosci. 47, 461–474 (2024). 10.1016/j.tins.2024.04.003

8 Zhang, P. et al. Senolytic therapy alleviates Aβ-associated oligodendrocyte progenitor cell senescence and cognitive deficits in an Alzheimer’s disease model. Nat. Neurosci. 22, 719–728 (2019). 10.1038/s41593-019-0372-9

9 Bussian, T. J. et al. Clearance of senescent glial cells prevents tau-dependent pathology and cognitive decline. Nature 562, 578–582 (2018). 10.1038/s41586-018-0543-y

10 Herdy, J. R. et al. Increased post-mitotic senescence in aged human neurons is a pathological feature of Alzheimer’s disease. Cell Stem Cell 29, 1637–1652.e1636 (2022). 10.1016/j.stem.2022.11.010

11 Pașca, S. P., et al. A nomenclature consensus for nervous system organoids and assembloids. Nature 609, 907–910 (2022). 10.1038/s41586-022-05219-6

12 Birtele, M., Lancaster, M. & Quadrato, G. Modelling human brain development and disease with organoids. Nat. Rev. Mol. Cell Biol. 26, 389–412 (2025). 10.1038/s41580-024-00804-1

13 Cherry, C. et al. Transfer learning in a biomaterial fibrosis model identifies in vivo senescence heterogeneity and contributions to vascularization and matrix production across species and diverse pathologies. Geroscience 45, 2559–2587 (2023). 10.1007/s11357-023-00785-7

14 Saul, D. et al. A new gene set identifies senescent cells and predicts senescence-associated pathways across tissues. Nat Commun 13, 4827 (2022). 10.1038/s41467-022-32552-1

15 Jones, R. C. et al. The Tabula Sapiens: A multiple-organ, single-cell transcriptomic atlas of humans. Science 376, eabl4896 (2022). 10.1126/science.abl4896

16 Hodge, R. D. et al. Conserved cell types with divergent features in human versus mouse cortex. Nature 573, 61–68 (2019). 10.1038/s41586-019-1506-7

17 Graham, V., Khudyakov, J., Ellis, P. & Pevny, L. SOX2 functions to maintain neural progenitor identity. Neuron 39, 749–765 (2003). 10.1016/s0896-6273(03)00497-5

18 Quinn, J. C., et al. Pax6 controls cerebral cortical cell number by regulating exit from the cell cycle and specifies cortical cell identity by a cell autonomous mechanism. Dev. Biol. 302, 50–65 (2007). 10.1016/j.ydbio.2006.08.035

19 Mahmud, F., Sarker, D. B., Jocelyn, J. A. & Sang, Q. A. Molecular and Cellular Effects of Microplastics and Nanoplastics: Focus on Inflammation and Senescence. Cells 13 (2024). 10.3390/cells13211788

20 Hwang, H. et al. Neurogranin, Encoded by the Schizophrenia Risk Gene NRGN, Bidirectionally Modulates Synaptic Plasticity via Calmodulin-Dependent Regulation of the Neuronal Phosphoproteome. Biol. Psychiatry 89, 256–269 (2021). 10.1016/j.biopsych.2020.07.014

21 Li, B., Woo, R. S., Mei, L. & Malinow, R. The neuregulin-1 receptor erbB4 controls glutamatergic synapse maturation and plasticity. Neuron 54, 583–597 (2007). 10.1016/j.neuron.2007.03.028

22 Liu, H. & Zhang, S. C. Specification of neuronal and glial subtypes from human pluripotent stem cells. Cell. Mol. Life Sci. 68, 3995–4008 (2011). 10.1007/s00018-011-0770-y

23 Stern, C. D. Neural induction: old problem, new findings, yet more questions. Development 132, 2007–2021 (2005). 10.1242/dev.01794

24 Leibovitz, Z., Lerman-Sagie, T. & Haddad, L. Fetal Brain Development: Regulating Processes and Related Malformations. Life (Basel) 12 (2022). 10.3390/life12060809

25 Nelson, C. A., 3rd & Gabard-Durnam, L. J. Early Adversity and Critical Periods: Neurodevelopmental Consequences of Violating the Expectable Environment. Trends Neurosci. 43, 133–143 (2020). 10.1016/j.tins.2020.01.002

26 Bourgeron, T. From the genetic architecture to synaptic plasticity in autism spectrum disorder. Nat. Rev. Neurosci. 16, 551–563 (2015). 10.1038/nrn3992

27 Birnbaum, R. & Weinberger, D. R. Genetic insights into the neurodevelopmental origins of schizophrenia. Nat. Rev. Neurosci. 18, 727–740 (2017). 10.1038/nrn.2017.125

28 Chow, K. H. & Abel, T. Neurodevelopmental origins of age-related neurodegenerative diseases. EBioMedicine 124, 106151 (2026). 10.1016/j.ebiom.2026.106151

29 Klein, A., Rhinn, M. & Keyes, W. M. Cellular senescence and developmental defects. Febs j 290, 1303–1313 (2023). 10.1111/febs.16731

30 Rhinn, M. et al. Aberrant induction of p19Arf-mediated cellular senescence contributes to neurodevelopmental defects. PLoS Biol. 20, e3001664 (2022). 10.1371/journal.pbio.3001664

31 Pietrogrande, G. et al. Valproic acid-induced teratogenicity is driven by senescence and prevented by Rapamycin in human spinal cord and animal models. Mol. Psychiatry 30, 986–998 (2025). 10.1038/s41380-024-02732-0

32 Heyer, D. B. & Meredith, R. M. Environmental toxicology: Sensitive periods of development and neurodevelopmental disorders. Neurotoxicology 58, 23–41 (2017). 10.1016/j.neuro.2016.10.017

33 Uhlhaas, P. J. & Singer, W. Neural synchrony in brain disorders: relevance for cognitive dysfunctions and pathophysiology. Neuron 52, 155–168 (2006). 10.1016/j.neuron.2006.09.020

34 OECD (2022). Global Plastics Outlook: Economic Drivers, Environmental Impacts and Policy Options. OECD Publishing. 10.1787/de747aef-en

35 OECD. Plastics Use - Estimations from 1990 to 2019. (2024). OECD Data Explorer

36 Sergiu Pasca, S.-J. Y. Feeder-free, Xeno-free Generation of Cortical Spheroids From Human Pluripotent Stem Cells. Protocol exchange (2018). 10.1038/protex.2018.123

37 Reumann, D. et al. In vitro modeling of the human dopaminergic system using spatially arranged ventral midbrain-striatum-cortex assembloids. Nat Methods 20, 2034–2047 (2023). 10.1038/s41592-023-02080-x

38 Abdulla, S. et al. CZ CELLxGENE Discover: a single-cell data platform for scalable exploration, analysis and modeling of aggregated data. Nucleic Acids Res. 53, D886–d900 (2025). 10.1093/nar/gkae1142

39 Wolf, F. A., Angerer, P. & Theis, F. J. SCANPY: large-scale single-cell gene expression data analysis. Genome Biol. 19, 15 (2018). 10.1186/s13059-017-1382-0

40 Hao, Y. et al. Dictionary learning for integrative, multimodal and scalable single-cell analysis. Nat. Biotechnol. 42, 293–304 (2024). 10.1038/s41587-023-01767-y

41 McInnes, L. UMAP: Uniform Manifold Approximation and Projection for Dimension Reduction. arXiv 1802.03426 (2018). <https://arxiv.org/abs/1802.03426v3>.

42 Marquez-Galera, A., de la Prida, L. M. & Lopez-Atalaya, J. P. A protocol to extract cell-type-specific signatures from differentially expressed genes in bulk-tissue RNA-seq. STAR Protoc 3, 101121 (2022). 10.1016/j.xpro.2022.101121

43 Cid, E. et al. Sublayer- and cell-type-specific neurodegenerative transcriptional trajectories in hippocampal sclerosis. Cell Rep. 35, 109229 (2021). 10.1016/j.celrep.2021.109229

44 Love, M. I., Huber, W. & Anders, S. Moderated estimation of fold change and dispersion for RNA-seq data with DESeq2. Genome Biol. 15, 550 (2014). 10.1186/s13059-014-0550-8

45 Langfelder, P., Zhang, B. & Horvath, S. Defining clusters from a hierarchical cluster tree: the Dynamic Tree Cut package for R. Bioinformatics 24, 719–720 (2008). 10.1093/bioinformatics/btm563

46 University of Colorado Boulder Research Computing (2021). PetaLibrary. University of Colorado Boulder. 10.25811/81nc-wv41

